# Cerebellar Microcircuits and Cortico–Cerebellar Cooperation Enable Robust Decisions

**DOI:** 10.64898/2026.04.26.720835

**Authors:** Yeyao Bao, Liao Yu, Liangfu Lv, Zhuoqing Yang, Yunliang Zang

## Abstract

The cerebellum is increasingly implicated in perceptual decision-making, yet how cerebellar circuits— without prominent recurrent excitation and slow reverberant loops—could support sparse evidence accumulation over behavioral timescales remains unclear. We present a biologically constrained modeling framework showing that cerebellar microcircuits can implement graded accumulation and competition without cortical-like excitatory recurrence. Type-II Purkinje-neuron excitability generates firing-rate hysteresis that prolongs the impact of brief inputs far beyond intrinsic membrane and synaptic time constants, enabling accumulation across long inter-event intervals. Purkinje neuron collateral inhibition produces competitive divergence and tunes temporal evidence weighting, revealing a trade-off between commitment and primacy bias. In a bidirectionally coupled cortico-cerebello-cortical model, cerebellar processing reduces primacy while cortical processing reduces indecision, improving robustness. Together, these results propose a mechanistic division of labor that positions the cerebellum as an active computational partner in perceptual decisions.

## INTRODUCTION

The cerebellum has long served as a model system for linking cellular physiology, canonical microcircuit architecture, and computation. Classic theories emphasized sensorimotor control and learning, leveraging stereotyped connectivity and well-characterized neuronal dynamics ^1-6^. Converging evidence from anatomy, electrophysiology, and imaging now implicates the cerebellum in cognitive functions, including working memory and decision-making ^7-14^. A central challenge is to move from phenomenology to mechanism: what computations does cerebellar circuitry contribute outside the motor domain, and through which biophysical and microcircuit substrates?

Perceptual decision-making provides a stringent test because it requires integrating noisy, sparse evidence over hundreds of milliseconds to seconds and converting that graded estimate into a categorical commitment ^9,10,15-18^. In cortical models, long-timescale integration is typically attributed to recurrent excitation, often stabilized by slow NMDA-mediated currents, and in some frameworks to multi-stage feedforward dynamics ^19-22^. Cerebellar cortex is organized differently: synaptic time constants are fast and the circuit does not prominently feature strong recurrent excitation ^1-3,23-29^, raising a mechanistic puzzle in light of recent causal and correlational results. Rodent evidence-accumulation tasks recruit cerebellar pathways ^7,9,10^, Purkinje-neuron perturbations impair choice accuracy, and Purkinje neurons exhibit cue-period bidirectional modulation during accumulation ^9,10^. Together, these observations suggest that cerebellar activity is not merely motoric or a passive readout of cortical decisions, but may participate in computations that shape choice.

Two questions follow. What cerebellum-specific neuronal and microcircuit mechanisms can expand the effective integration window to support sparse, stochastic evidence accumulation without relying on cortical-style recurrent excitation or multi-stage feedforward dynamics? If both cortex and cerebellum influence decisions, what computational advantage is gained by their interaction, and what division of labor makes the coupled system more robust than either module alone?

Here we develop a biologically constrained modeling framework grounded in rodent evidence-accumulation paradigms in which sensory evidence arrives as randomly timed pulses ^9,10,18^. We first analyze a cerebellar microcircuit model to identify intrinsic and local mechanisms sufficient for accumulation and competition. We then embed this circuit in coupled cortico-cerebellar architectures to ask how bidirectional interactions alter decision performance. By comparing cerebellar-only, cortico-cerebellar, cortico-cerebello-cortical, and cortex-only models, we isolate the roles of Purkinje-neuron excitability, Purkinje collateral inhibition, granule layer sparsification, and cortico-cerebellar coupling.

We identify an intrinsic Purkinje-neuron mechanism that extends the effect of brief inputs to support low-frequency accumulation, and a local inhibitory mechanism that tunes temporal evidence weighting, revealing a trade-off between commitment and primacy. We further show that granule layer sparsification improves discrimination when evidence streams are correlated, and that closed-loop cortico-cerebellar coupling yields a division of labor that improves robustness across stimulus statistics and timing variability. Finally, we derive testable predictions linking Purkinje collateral inhibition and Golgi-cell-mediated sparsification to measurable changes in temporal kernels and choice behavior, and predicting complementary cortical versus cerebellar signatures across task regimes.

## RESULTS

### Cerebellar circuits contribute to evidence accumulation in decision-making

To examine how cerebellar circuitry could support perceptual decisions, we modeled rodent evidence-accumulation paradigms that probe decisions under temporal uncertainty (Figure 1A). We focus on the fixed-time task, in which stochastic air puffs are delivered to left and right whiskers during a fixed cue epoch, followed by a delay and a choice report. Trained animals achieve high accuracy, and Purkinje neurons exhibit cue-period ramping or bidirectional modulation of firing rates ^9,10^. We also simulated the reaction-time task (Figure 7 and S5), in which cue duration is variable and animals respond as soon as they infer which side has the higher Poisson input rate ^35^.

**Figure 1.**
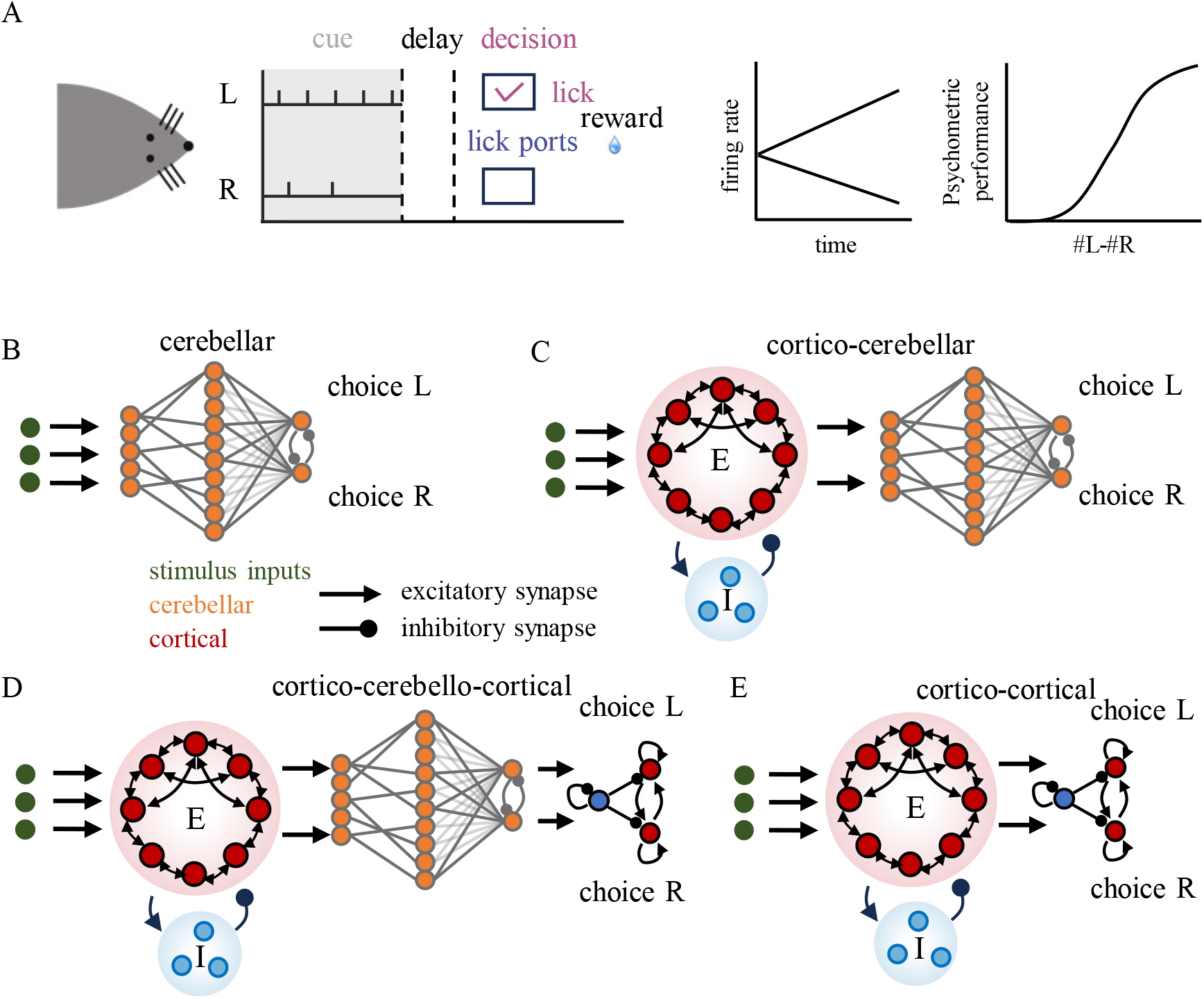
Experimental paradigms and alternative circuit models. (A) Rodent evidence-accumulation task. A fixed cue (evidence) period is followed by a delay and a choice report (lick to the side with more stimuli) ^9,10,18^. Schematics of Purkinje-neuron firing rates (middle) and psychometric performance (right) are shown. (B) Cerebellar-only model driven by sensory-triggered mossy-fiber spike bursts ^30^. Purkinje neurons are coupled by inhibitory axon collaterals ^31-33^. (C) Cortico-cerebellar model in which a cortical continuous-attractor module preprocesses sensory inputs before they enter the cerebellum ^34^. (D) Cortico-cerebello-cortical model extending (C) with a cortical decision module that integrates evidence and generates categorical choices ^19^. (E) Cortico-cortical model, matched to (D) but without the cerebellar module. The two cortical modules differ primarily in connectivity and parameter settings; schematics emphasize their distinct functions. See Method Details.

Across models, we do not explicitly simulate motor execution. Instead, a “decision” is registered when the firing-rate difference between two competing populations exceeds a fixed threshold ^36^, allowing a common readout across architectures.

We compared four circuit architectures (Figure 1B-E). The pure cerebellar model isolates intrinsic and local circuit mechanisms for accumulation and competition, with mossy fibers providing discrete spiking bursts motivated by bouton recordings ^30^. The cortico–cerebellar model adds a cortical continuous-attractor module that generates heterogeneous, temporally dispersed mossy-fiber activity consistent with imaging data ^34,37^. The cortico-cerebello-cortical model closes the loop by adding a cortical decision network ^19^ downstream of the cerebellum ^38-40^, enabling us to test how coupling two partially redundant decision-capable modules changes performance. A cortex-only (cortico–cortical) model serves as a reference for quantifying the computational advantages conferred by cerebellar involvement compared to cortical-only processing.

Together, these models provide a controlled framework to ask (i) how cerebellar microcircuits can integrate sparse evidence despite fast synapses and short membrane time constants, and (ii) what computational advantage emerges from cortico-cerebellar coupling.

### Type-II excitability enables low-frequency accumulation in Purkinje neurons

We first asked whether a cerebellar circuit can integrate sparse sensory events in the absence of cortical-like mechanisms. In the pure cerebellar model, left- and right-selective mossy-fiber populations deliver brief spike bursts representing discrete sensory events (Figure 2). Granule cell→Purkinje neuron synapses were modeled as AMPA conductances with a 2 ms decay time constant, and Purkinje neuron model had an intrinsic integration time constant of ∼32 ms ^24,41,42^. Inhibitory axonal collaterals between Purkinje neurons were modeled as GABA conductances ^31-33^.

**Figure 2.**
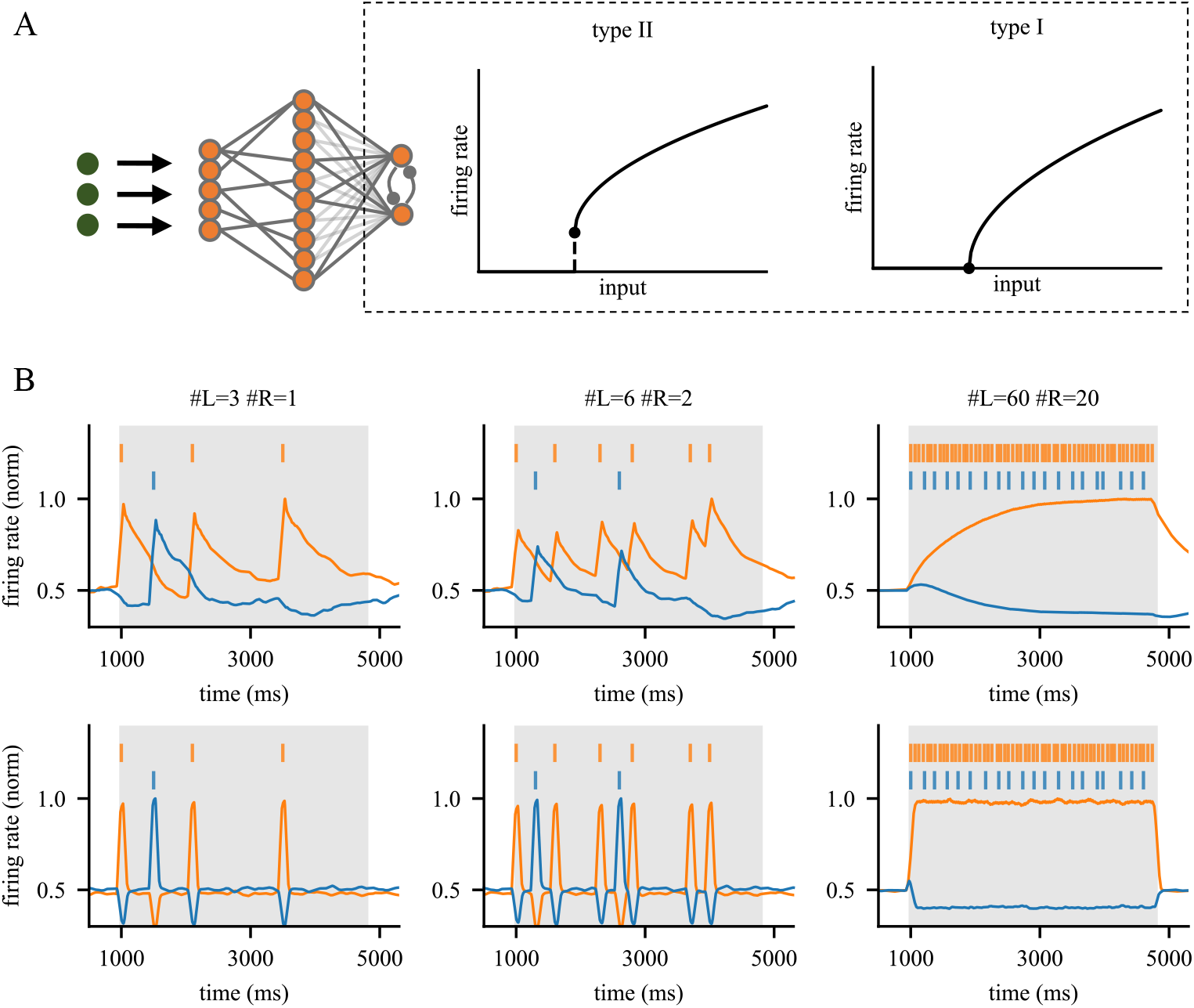
Type-II excitability enables low-frequency evidence integration in Purkinje neurons. (A) Cerebellar circuit model with Purkinje neurons implemented with type-II or type-I excitability. Type-I and type-II models exhibit continuous versus discontinuous f-I relationships, respectively. (B) Purkinje-neuron firing-rate dynamics under increasing event rates (left to right). Vertical lines mark stimulus events; traces show firing rates of left- and right-selective populations.Top and bottom panels correspond to type-II and type-I excitability, respectively.

In a biophysically grounded type-II Purkinje-neuron model, population activity ramps during the cue period: the population receiving stronger evidence increases firing while the competing population decreases firing. Notably, ramping persists even when events are extremely sparse (0.8 Hz; inter-event interval 1250 ms), far exceeding intrinsic membrane and synaptic time constants.

Replacing type-II with type-I excitability abolishes low-frequency accumulation. Type-I models only show effective accumulation at high event rates when inter-event intervals are shorter than intrinsic time constants; at experimentally tested rates (2.5–5.0 Hz ^9,10,18^), stimulus-evoked responses completely recover before the next event, and activity fails to persist after cue period required by the fixed-time task for decision categorization ^9,10,18^.

Purkinje neuron collateral inhibition generates opposing population trajectories during accumulation; removing this inhibition eliminates competitive divergence (Figure S1). These results link sparse-event accumulation to type-II excitability and motivate a mechanistic question: which feature of type-II dynamics extends the effective integration window?

### Firing‐rate hysteresis extends the temporal window of neuronal integration

To identify the mechanism, we analyzed the near-threshold input–output dynamics of the type-I and type-II models. Type-II models exhibit firing-rate hysteresis during slow current ramps (Figure 3A), consistent with experimentally observed distinct “up” and “down” firing regimes ^43,44^. In contrast, type-I models show a single-valued, history-independent firing-rate curve.

**Figure 3.**
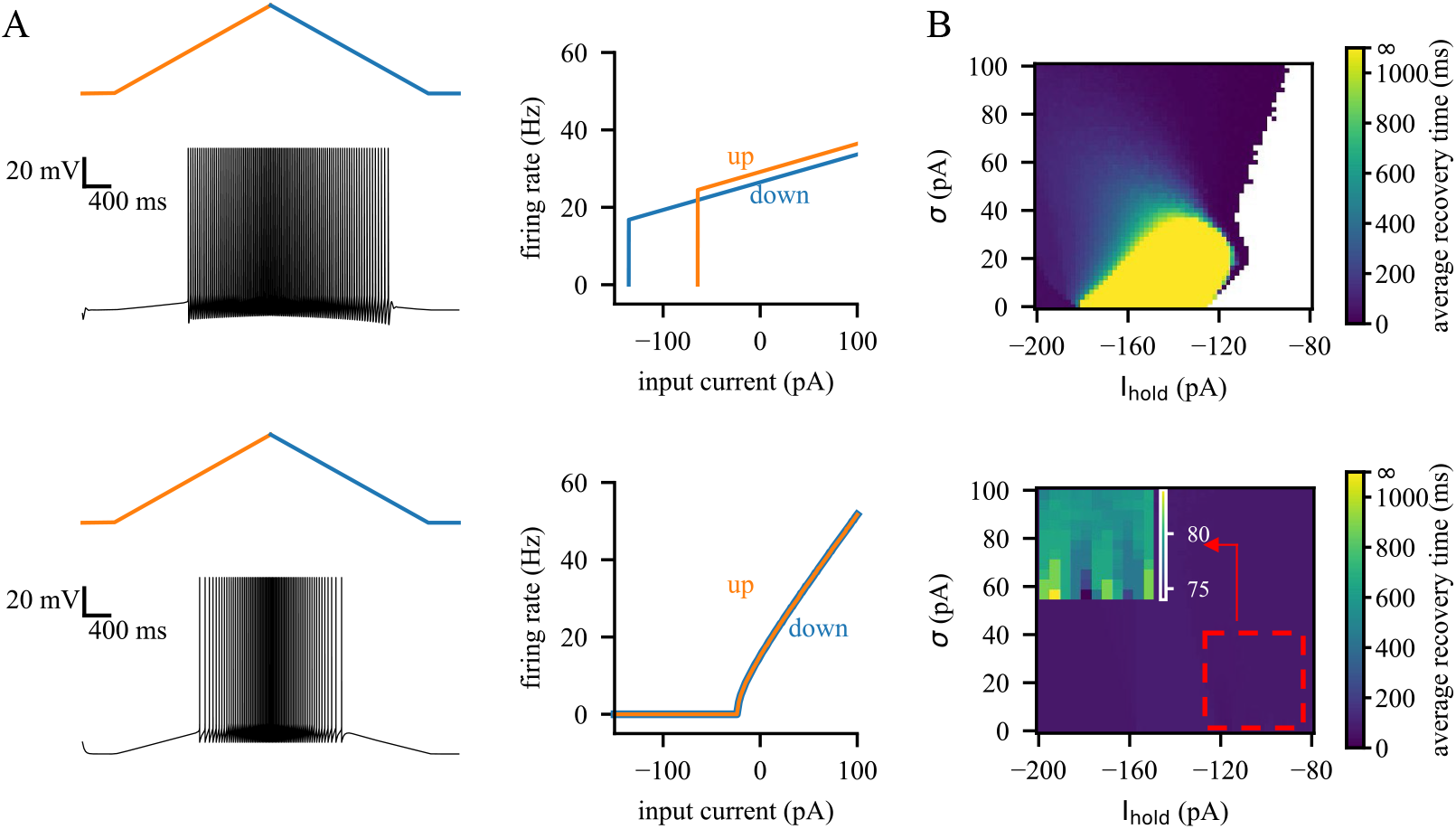
Hysteresis in Purkinje-neuron firing extends the integration window. (A) Firing-rate hysteresis in type-II models during slow current ramps (top) compared to the continuous response of type-I models (bottom). (B) Recovery time constant as a function of I_hold_ and noise variance σ. The white region indicates parameter regimes in which neurons are entirely in the firing regime at baseline and never return to silence.

We quantified the functional consequence of hysteresis by applying a holding current (*I*_*hold*_) and Gaussian noise (variance *σ*), and then measuring recovery time following a single transient stimulus (Figure 3B). In the type-II model, once a stimulus drives a neuron into the firing regime, returning to silence requires a larger opposing drive, producing slow recovery at the population level. Recovery time constants frequently exceed 1 sec across a broad parameter range, orders of magnitude longer than the tens-of-milliseconds recovery observed in the type-I model.

Type-I models could not reproduce this slow recovery because their upward and downward thresholds are symmetric (Figure S2). Thus, firing-rate hysteresis provides a single-cell mechanism that prolongs the impact of brief inputs and supports sparse-event accumulation.

### Cortical preprocessing enhances cerebellar integration

Pontine mossy fibers can exhibit heterogeneous, delayed responses to a discrete sensory event ^34,37^. To capture this, we implemented a cortical continuous-attractor module that transforms each discrete stimulus into a temporally dispersed population response, producing mossy-fiber spike trains with variable latencies (Figure 4A).

**Figure 4.**
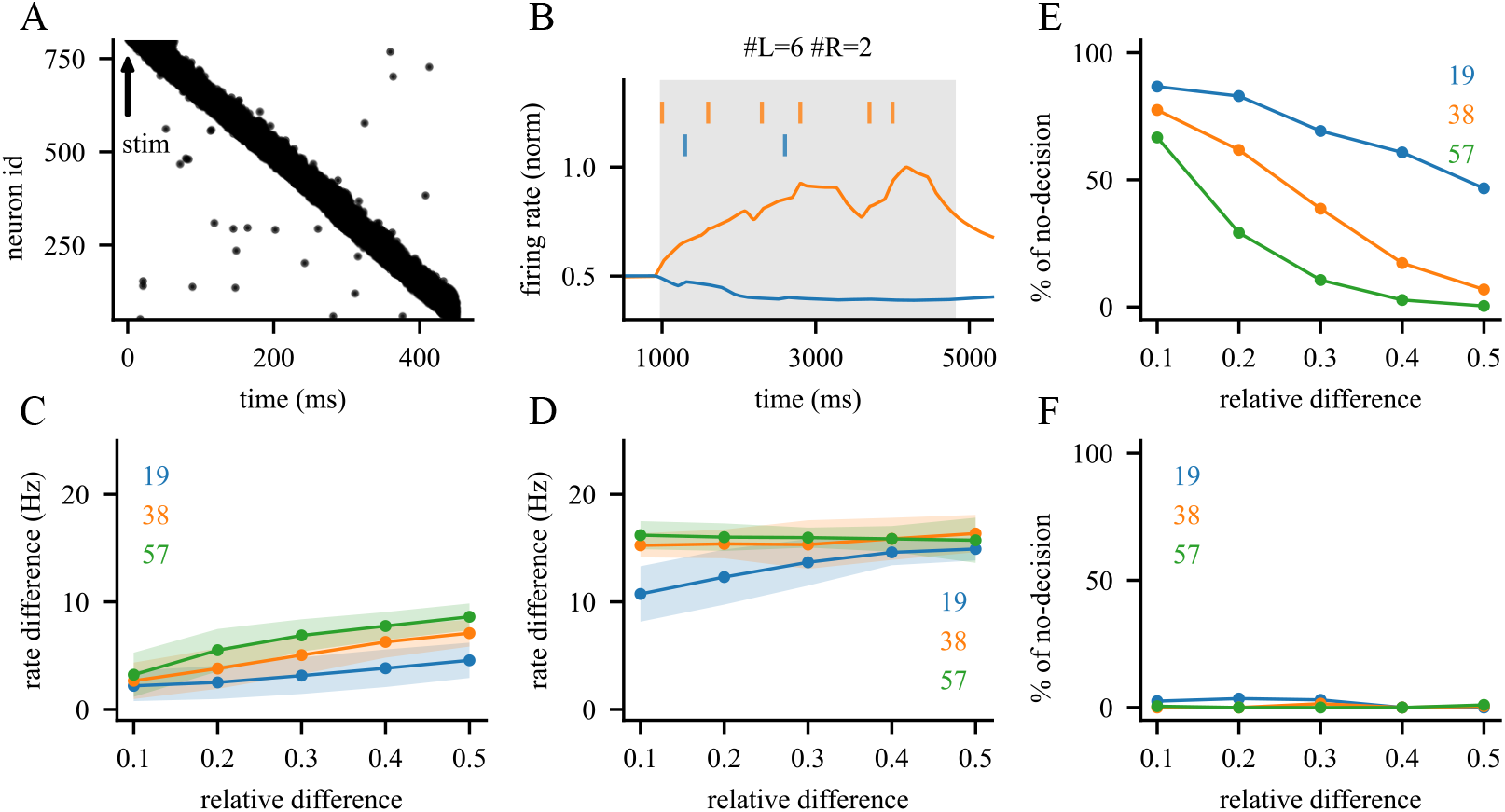
Cortical preprocessing enhances cerebellar evidence accumulation. (A) Example mossy-fiber spike trains with heterogeneous latencies generated by the cortical continuous attractor module. (B) Purkinje-neuron ramping under low-frequency evidence. (C–D) Decision readiness (cue-end firing-rate difference) for spike bursts (C) versus cortical-preprocessed inputs (D). (E–F) Proportion of undecided trials as a function of relative evidence difference for direct bursts (E) versus preprocessed inputs (F). Colored numbers indicate total evidence across both sides.

When these preprocessed inputs drive the cerebellar circuit, Purkinje-neuron ramping becomes more pronounced under low event rates (compare Figure 4B with Figure 2B). Cortical preprocessing increases the firing-rate separation between competing Purkinje neuron populations at cue end (Figure 4C–D), thereby increasing “decision readiness” under a fixed-threshold readout.

Consistent with increased cue-end separation, cortical preprocessing reduces the fraction of undecided trials, particularly at low event rates (Figure 4E–F). These results suggest a division of labor: cortical preprocessing distributes evidence over time, while cerebellar hysteresis integrates and stabilizes it.

### Purkinje-neuron mutual inhibition tunes temporal evidence weighting

We next examined how local mutual inhibition mediated by Purkinje neuron axon collaterals shapes competition and temporal weighting. When inhibitory conductance exceeds ∼18 nS, network activity becomes unstable and enters self-sustained bistable states even without sensory drive (Figure S3) ^45^. Within the stable regime, psychometric performance varies non-monotonically with inhibition strength (Figure 5A).

**Figure 5.**
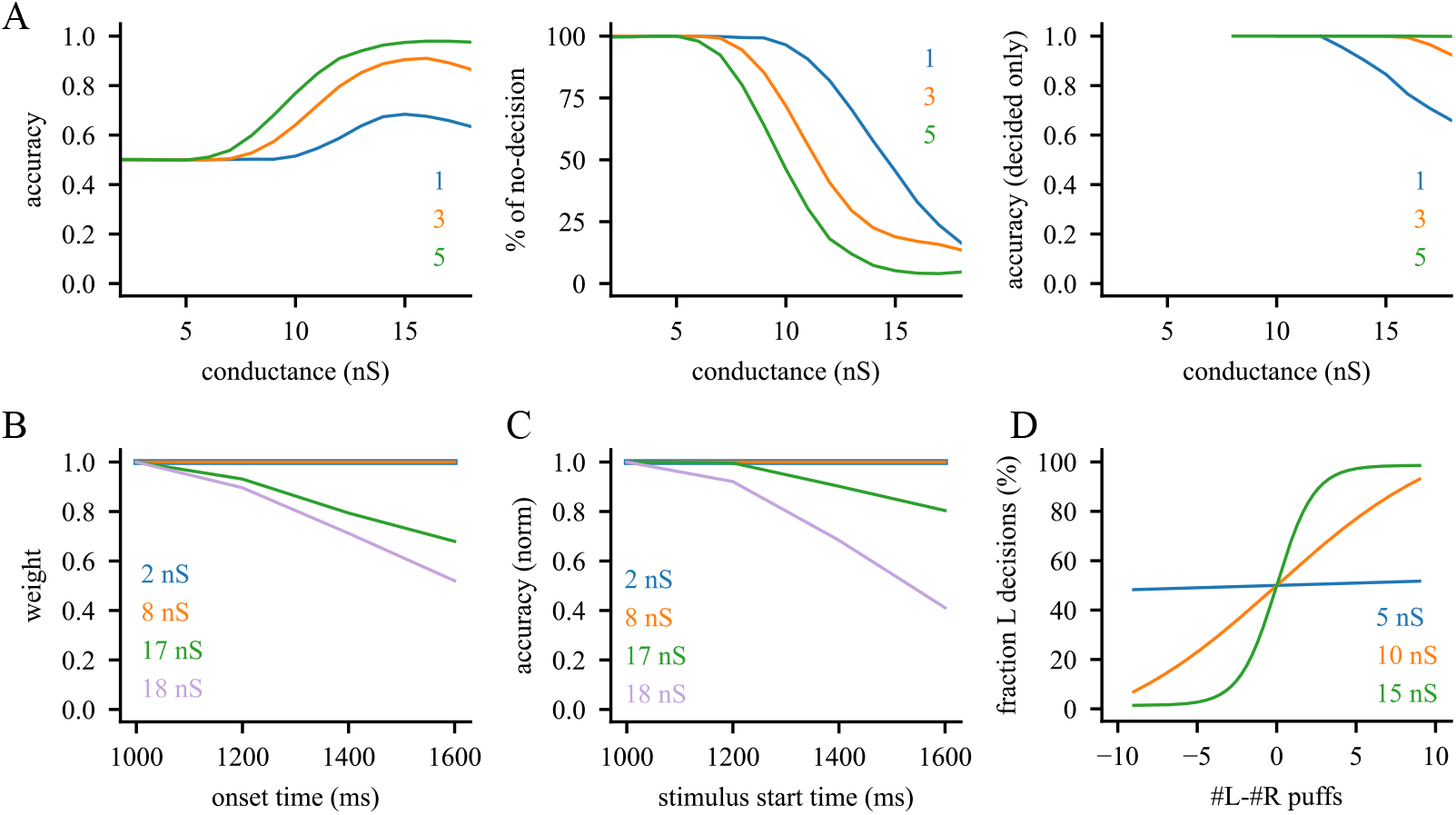
Mutual inhibition shapes accumulation and temporal weighting. (A) Psychometric curves versus mutual inhibition conductance between Purkinje neurons. Left: accuracy including undecided trials. Middle: undecided fraction. Right: accuracy excluding undecided trials. Colors indicate evidence-count differences. (B) Temporal weighting (psychophysical kernel) of a pulse as a function of pulse time. (C) Accuracy versus preferred-stream onset time (non-preferred onset fixed). (D) Psychometric curves versus relative evidence difference. Colored numbers in A indicate the difference between evidence between the two sides. Colors in B–D indicate inhibition strength.

Weak inhibition increases trial-to-trial variability in cue-end separation and increases the fraction of undecided trials, reducing overall accuracy (Figure 5A, left and middle). Conditioning on committed trials reveals that weak inhibition could improve accuracy among decided trials (Figure 5A, right), suggesting a trade-off between commitment rate and integration quality.

To examine how inhibition strength regulates integration quality, we assessed temporal weighting using a psychophysical pulse paradigm ^46^: transient pulses delivered at variable latencies on top of a constant background reveal a primacy-like weighting profile (Figure 5B) ^47,48^. Weak inhibition attenuates primacy, producing more uniform weighting across time. This reduced temporal bias improves robustness to stochastic evidence timing: when the non-preferred stream begins at a fixed time and the preferred stream is delayed, strong inhibition causes pronounced accuracy loss whereas weak inhibition preserves performance (Figure 5C). However, weak inhibition yields shallower psychometric curves because more trials fail to reach the decision threshold (Figure 5D).

These results identify local mutual inhibition among Purkinje neurons as a control parameter that trades off commitment (fewer undecided trials) against temporal bias (stronger primacy).

### Cortico-cerebello-cortical coupling balances primacy bias and indecision

Motivated by higher-order cerebello-cortical feedback pathways ^7,49-52^, we next asked whether adding a downstream cortical decision module can resolve the bias–indecision trade-off. We therefore coupled the cerebellar circuit to a cortical decision module ^19^, yielding a cortico-cerebello-cortical model, and also compared it to a cortex-only model with matched cortical dynamics.

Adding the cortical module reduces undecided trials and improves accuracy in the weak-inhibition regime relative to the cortico-cerebellar model, although performance decreases slightly at high Purkinje neuron inhibition range (Figure 6A). Across inhibition values, stronger Purkinje neuron inhibition increases cue-end separation (Figure 6B) but also increases primacy bias (Figure 6C-D). At extremely weak inhibition, cortical attractor dynamics dominates temporal weighting, producing a primacy profile similar to the cortex-only model.

**Figure 6.**
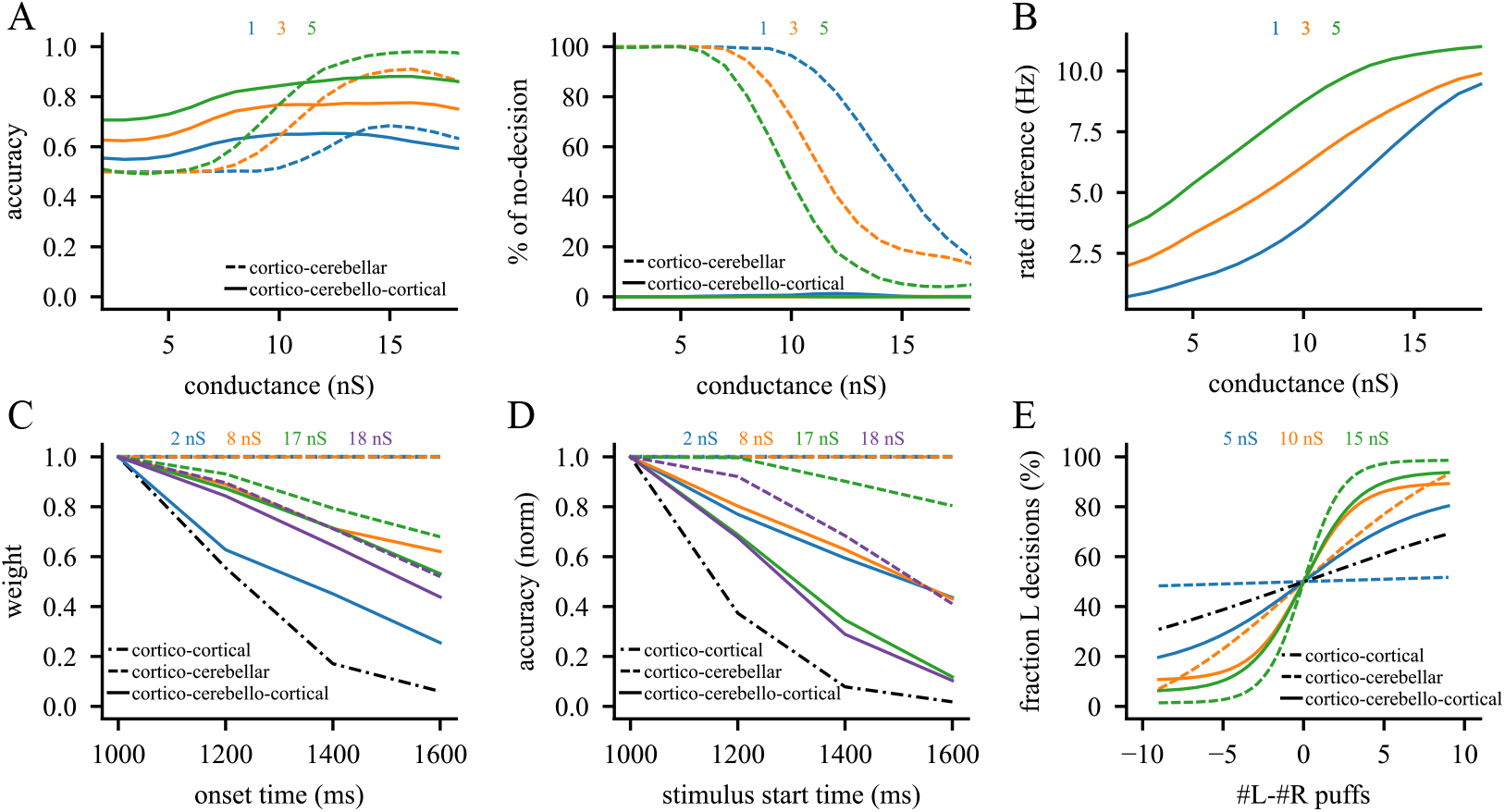
Closed-loop coupling balances temporal bias and commitment. (A) Psychometric curves versus Purkinje neuron mutual inhibition conductance. Left: accuracy including undecided trials. Right: undecided fraction. (B) Cue-end decision readiness versus mutual inhibition conductance. (C) Temporal weighting versus pulse time. (D) Accuracy versus preferred-stream onset time (non-preferred fixed). (E) Psychometric curves versus relative evidence difference. In A-B, colors indicate evidence-count differences. In C-E, colors indicate Purkinje neuron inhibition strength; solid: cortico-cerebello-cortical; dashed: cortico-cerebellar; dash-dotted: cortex-only.

**Figure 7.**
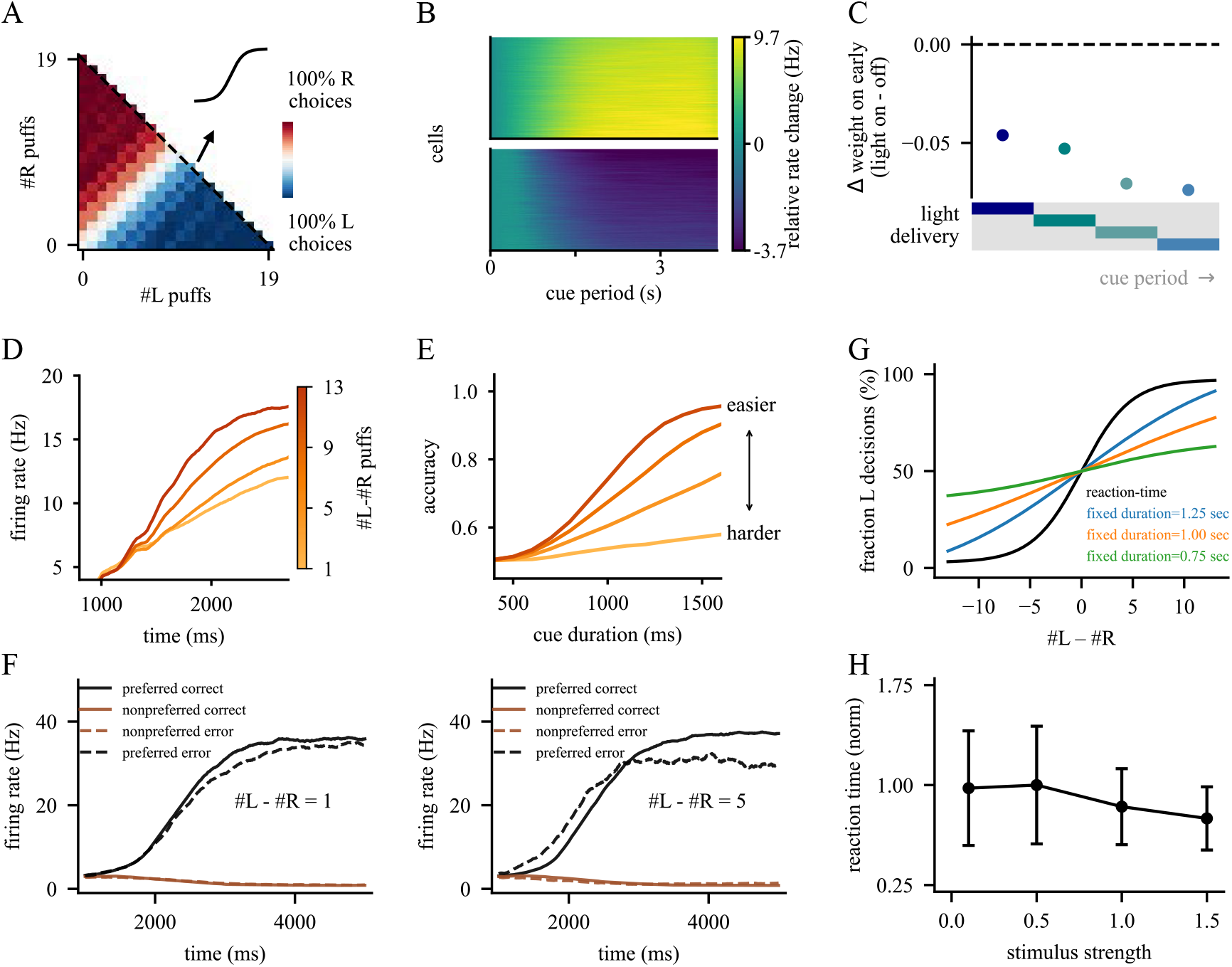
Evidence-accumulation dynamics captured by the cortico-cerebello-cortical model. (A) Choice probability as a function of left and right puff counts. (B) Cue-period activity of winning (top) and losing (bottom) Purkinje neuron populations. (C) Weight change following cue-period Purkinje neuron perturbations applied at different times. (D) Winning-population Purkinje neuron trajectories across evidence differences. (E) Psychometric curves across cue durations and difficulty levels. (F) Population trajectories for correct and error trials across stimulus differences. (G) Psychometric curves comparing different fixed cue durations with the reaction-time paradigm. (H) Mean reaction time changes versus stimulus strength. Panels A-F: fixed-time; panel H: reaction-time; panel G: both.

Across the stable conductance range, temporal bias ranks consistently: the cortico-cerebellar model shows the weakest primacy, the cortico-cerebello-cortical model is intermediate, and the cortex-only model shows the strongest primacy (Figure 6C-D). Consistent with this complementarity, the coupled model achieves high performance across a broader range of Purkinje neuron inhibition strengths than either module alone (Figure 6E).

These results support a computational division of labor: cerebellar dynamics reduce temporal bias during evidence integration, whereas the cortical module promotes categorical commitment and reduces indecision. Example circuit dynamics illustrating these roles are shown in Figure S4.

### Cortico-cerebello-cortical model reproduces behavioral and neuronal signatures

We next asked whether the cortico-cerebello-cortical model reproduces hallmark behavioral and neural signatures of evidence accumulation. In the fixed-time task, the model reproduces choice probabilities as a function of left and right puff counts (Figure 7A) ^9^. Purkinje neuron activity ramps during the cue period, with the winning population increasing and the losing population decreasing (Figure 7B) ^9^. Cue-period perturbations of Purkinje neuron activity reduces accuracy, with later perturbations producing larger impairments, consistent with experimentally observed evidence discounting (Figure 7C) ^9^. The slope of the winning trajectory scales with evidence strength (Figure 7D) ^9^, and accuracy increases with cue duration for difficult discriminations while saturating for easier ones (Figure 7E) ^18^. Population trajectories capture correct versus error dynamics: at small evidence differences, trajectories are similar across trial types, whereas larger differences produce stronger divergence with reduced slopes on error trials (Figure 7F) ^51,53^.

In the reaction-time task, decisions are registered once the firing-rate difference crosses a threshold ^35,36^. The model reproduces improved performance relative to short fixed cue durations (Figure 7G) ^18,51^, and mean reaction time decreases with increasing stimulus strength (Figure 7H) ^35^. Notably, these experimental signatures were already reproduced by the cortico-cerebellar model (Figure S5).

### Cerebellar pattern separation supports decisions under overlapping inputs

Finally, we tested whether cerebellar computations provide an advantage when sensory evidence is encoded by overlapping populations, a common regime in biological circuits ^35,54,55^ but not explicitly explored in previous models ^19,56,57^. We introduced mixed representations at the mossy-fiber level by assigning subsets of mossy fibers to respond preferentially to left, right, or both streams (Figure 8A).

**Figure 8.**
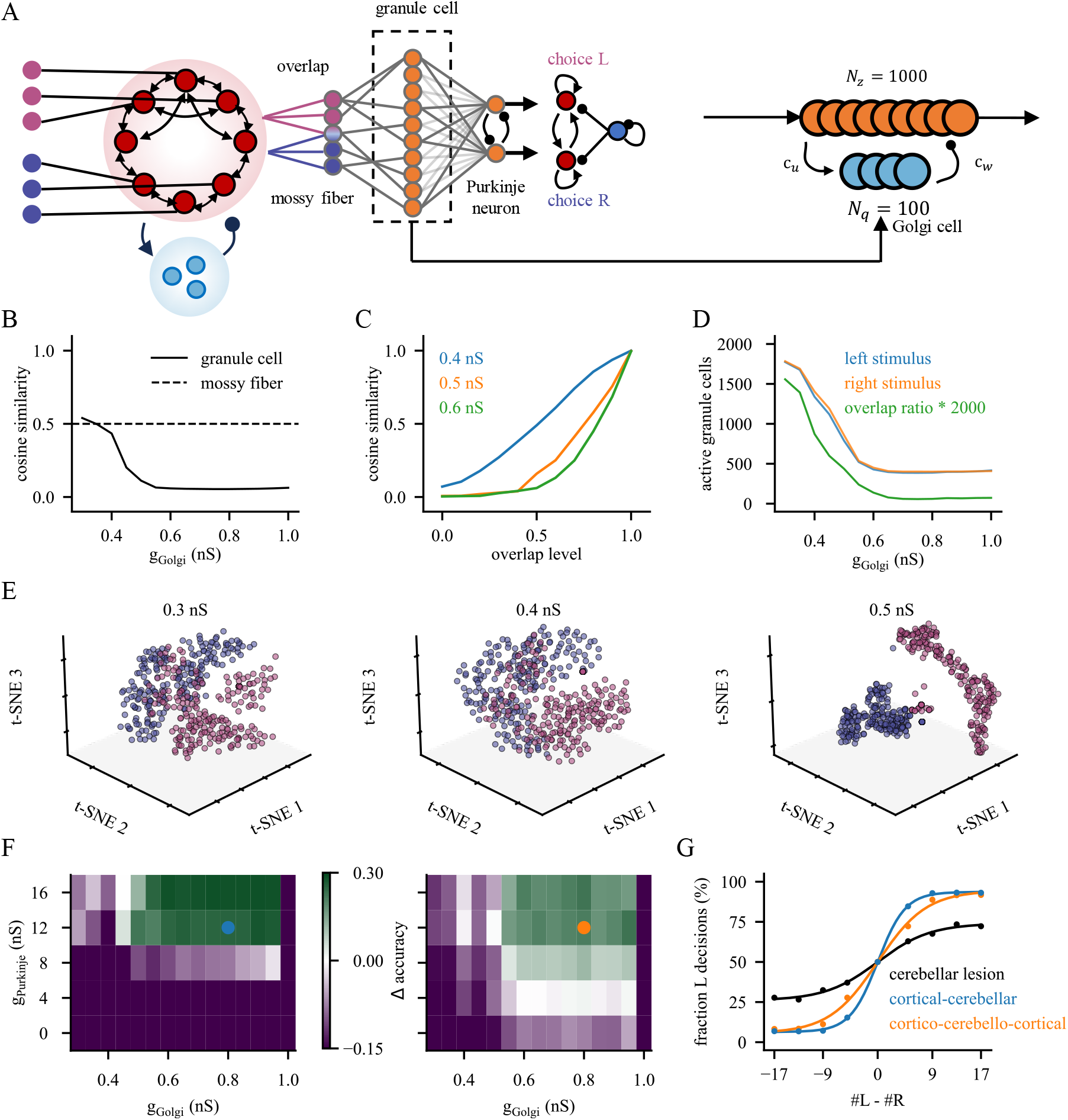
Cerebellar pattern separation improves decision reliability under overlapping inputs. (A) Mixed mossy-fiber encoding of left and right evidence; granule-cell sparsity is controlled by Golgi cell feedback inhibition. (B) Granule-cell response similarity versus Golgi cell conductance (dashed: mossy-fiber input similarity). (C) Granule-cell response similarity versus the fraction of mixed mossy fibers; colors indicate Golgi-cell conductance. (D) Number of active granule cells and response overlap versus Golgi-cell conductance. (E) t-SNE visualization of granule-cell population responses as Golgi cell inhibition increases. (F) Accuracy gains relative to the cortex-only model across Golgi-cell and Purkinje-neuron inhibition strengths (left: cortico-cerebellar minus cortex-only; right: cortico-cerebello-cortical minus cortex-only). (G) Psychometric performance versus stimulus strength across models. “Cerebellar lesion” corresponds to cortico-cortical model; the other curves correspond to parameter settings indicated in (F).

In the cerebellar cortex, granule cells receive sparse mossy-fiber inputs and are regulated by Golgi-cell inhibition. Increasing Golgi cell inhibition produces sparser granule cell activation, reduces response similarity relative to mossy-fiber inputs, and enhances pattern separation (Figure 8B-D). Consistent with this, t-SNE embeddings of granule cell population responses show progressively improved separation of left versus right patterns with stronger Golgi cell inhibition (Figure 8E). Improved separation increases discrimination and accumulation performance when mossy-fiber inputs are overlapped (Figure 8F-G).

Across a broad range of Golgi cell and Purkinje neuron inhibition strengths, the cortico-cerebellar model outperforms the cortex-only model. The full cortico-cerebello-cortical model maintains robust performance across a wider parameter range, though with smaller peak gains. These conclusions hold across overlap ratios (Figure S6). By contrast, the cortex-only model does not reach ceiling accuracy even at high evidence differences, consistent with experimental observations ^9^.

## DISCUSSION

This study proposes a mechanistic account of how cerebellar microcircuits can contribute directly to perceptual decision-making. Across model variants, three principles emerge. First, intrinsic Purkinje-neuron dynamics can provide a slow, history-dependent state variable that supports sparse-event accumulation despite fast synapses and short membrane time constants. Second, cerebellar local inhibition shapes competitive divergence and tunes the temporal weighting of evidence. Third, bidirectional cortico-cerebello-cortical coupling can resolve a trade-off between temporal bias and categorical commitment, improving robustness across stimulus statistics, temporal uncertainty, and correlated inputs.

A central challenge for accumulation under naturalistic conditions is that evidence often arrives as sparse, stochastic events whose inter-event intervals exceed intrinsic neuronal and synaptic integration time constants—typically tens of milliseconds ^24,41,42,58^. Distinct from cortical strategies ^19-21,59^, the cerebellum does not prominently feature strong recurrent excitation, motivating the question of how it could support integration. Our work identifies firing-rate hysteresis arising from type-II Purkinje-neuron excitability as a candidate solution ^43,44^: brief inputs can trigger recovery times on the order of seconds, effectively extending the temporal footprint of each event and enabling accumulation across long inter-event intervals. More broadly, this suggests a neuronal strategy for expanding integration windows that is distinct from cortical-style reverberation and may generalize to other circuits whose principal neurons exhibit type-II excitability ^60,61^.

Beyond intrinsic dynamics, our model assigns a functional role to Purkinje neuron axon collaterals. While these connections have been studied in the context of synchrony and development ^31-33,62^, their computational contribution remains unclear. In our framework, local mutual inhibition among Purkinje neurons generates competitive divergence between populations representing alternative evidence streams and tunes temporal weighting. Increasing collateral inhibition strengthens cue-end separation and reduces indecision but also increases primacy bias and can destabilize dynamics at high conductance (Figure S3). We therefore interpret Purkinje neuron collateral inhibition as a microcircuit mechanism that shapes the psychophysical kernel, trading off commitment against temporal bias.

The cerebellum also operates within distributed loops. Mossy-fiber inputs reflect transformations performed by upstream forebrain circuits, and imaging suggests that pontine inputs can be temporally dispersed and heterogeneous ^34,37^. In our model, a cortical preprocessing module distributes each sensory event over time, increasing cue-end separation and reducing undecided trials, particularly under sparse evidence; this facilitation persists in the closed-loop cortico-cerebello-cortical model (Figure S7). These results support a complementary interaction in which cortical dynamics diversify and temporally extend evidence representations, while cerebellar hysteresis integrates and stabilizes them.

Recent work has proposed that sequential, rather than persistent, activity can encode accumulated evidence through spatiotemporally tiled population dynamics ^63^. Our framework is compatible with this broader idea in that temporally structured input patterns—whether generated by a continuous attractor as modeled here or by alternative sequential-coding mechanisms—can be converted into stable, graded decision variables by cerebellar microcircuit dynamics. An important next step will be to test how different upstream coding schemes for evidence (persistent, sequential, or mixed) interact with cerebellar integration mechanisms to shape psychophysical kernels and neural trajectories.

At the systems level, we find that bidirectional cortico-cerebellar coupling can improve robustness by combining complementary failure modes. In the coupled cortico-cerebello-cortical model, cerebellar dynamics reduce primacy bias relative to cortex-only integration, whereas the cortical module reduces indecision near the choice boundary. This division of labor yields strong performance across a broader parameter range than either module alone, consistent with the idea that distributed circuits can implement similar computations through distinct mechanisms and thereby increase robustness to changes in input statistics and circuit state.

Finally, our results highlight a cerebellar advantage when evidence representations are overlapping or correlated. Many decision models assume distinct populations for each alternative ^19,21,56,57^, but real sensory pathways often carry mixed signals: mossy fibers and other precerebellar nuclei can receive bilateral and convergent inputs, and in modalities such as audition, left/right evidence cannot be perfectly segregated ^35,54,55^. In this regime, correlated inputs can degrade discrimination in cortex-only models. In contrast, Golgi cell-controlled sparsification in the granule layer decorrelates mixed mossy-fiber patterns and improves separability, extending classical expansion-recoding ideas ^64,65^ to decision computations. This suggests a general role for cerebellar pattern separation in cognitive settings where evidence streams are ambiguous or not cleanly partitioned.

### Limitations and future directions

Our models abstract away several biological features, including explicit thalamic relay dynamics and contributions from basal ganglia and other regions ^7,66,67^, and use a threshold-based readout rather than an explicit motor implementation ^36,68^. These simplifications allow us to isolate cerebellar microcircuit mechanisms, but future work should embed the present framework within more complete architectures and test whether the proposed division of labor persists under richer motor and state-dependent constraints. Core predictions can be tested experimentally: manipulating Purkinje neuron axon-collateral inhibition should bidirectionally modulate primacy in the psychophysical kernel and commitment rate; Golgi-cell perturbations ^69^ should preferentially impair performance when left/right evidence streams are correlated; and simultaneous cerebellar–cortical recordings during decision-making tasks should reveal cerebellar signatures associated with reduced temporal bias alongside cortical dynamics associated with categorical commitment.

## METHOD DETAILS

The model code will be made public once the manuscript is accepted.

### Cortical Continuous Attractor Module

The continuous attractor model serves to preprocess sensory inputs, transforming discrete stimuli into heterogeneous mossy fiber activities with variable onset times ^34,37^. The network consists of two neuronal populations: an excitatory (E) population of *N*_*E*_ (*N*_*E*_ = 1600) pyramidal neurons and an inhibitory (I) population of *N*_*I*_ (*N*_*I*_ = 400) interneurons ^70^. The excitatory neurons are functionally arranged along a ring structure representing a continuous feature space.

Both pyramidal neurons and interneurons are modeled as conductance-based leaky integrate-and-fire (LIF) model. The membrane potential *V*_*i*_ (*t*) of neuron *i* evolves according to:

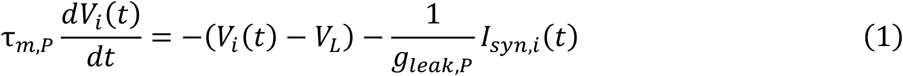

Where *P*, either E or I, denotes the population type. Here, *τ*_*m,P*_ is the membrane time constant, *V*_*L*_ is the leak reversal potential, and *g*_*leak,P*_ is the leak conductance. When *V*_*i*_ (*t*) exceeds the threshold *V*_*th*_, the neuron emits a spike, after which the potential is reset to *V*_*reset*_ and held for a refractory period*τ*_*ref,P*_. *I*_*syn,i*_ (*t*) represents the total synaptic current received by the neuron. Here, *τ*_*m,E*_ = 20 *ms, τ*_*m,I*_ = 10 *ms, V*_*L*_ = −70 *mV, g*_*leak,E*_ = 25 *nS, g*_*leak,I*_ = 20 *nS, V*_*th*_ = −50 *mV, V*_*reset*_ = −55 *mV, τ*_*ref,E*_ = 2 *ms, τ*_*ref,I*_ = 1 *ms*.

The network features all-to-all recurrent connectivity. Inhibitory projections are spatially uniform and independent of target tuning. Recurrent excitatory currents are mediated by AMPA and NMDA receptors, while inhibitory currents are mediated by GABA receptors. Neurons also receive external excitatory inputs through AMPA receptors, representing both sensory drive and background noise.

For an excitatory neuron:

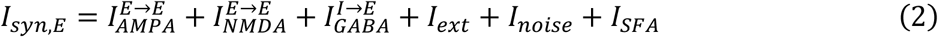

For an inhibitory neuron:

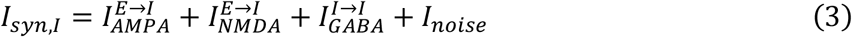

In Equations 2-3, the first three terms describe recurrent projections from the excitatory and inhibitory populations; *I*_*ext*_ denotes task-related external input signals, *I*_*noise*_ represents stochastic background noise, and *I*_*SFA*_ denotes spike-frequency–adaptation currents (included only in excitatory neurons).

### Synaptic Dynamics

Synaptic currents mediated by AMPA and GABA receptors follow:

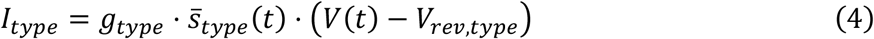

where *type* either represents AMPA or GABA, and *V*_*rev,type*_ is the corresponding reversal potential.

NMDA currents include a voltage‐dependent magnesium block:

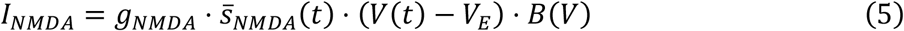

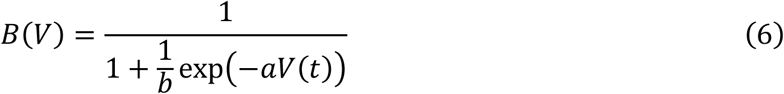

Spike‐frequency adaptation in excitatory neurons is modeled as:

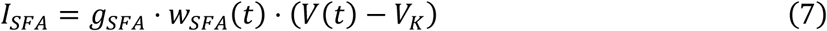

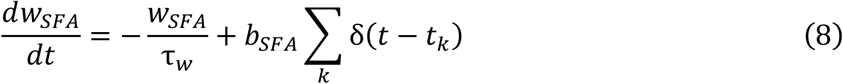

Here, *g*_*AMPA,E*_ = 150 *nS*, *g*_*AMPA,I*_ = 300 *nS*, *g*_*NMDA,E*_ = 400 *nS*, *g*_*NMDA,I*_ = 100 *nS*, *V*_*rev,E*_ = 0 *mV*, *V*_*rev,I*_ = −70 *mV*, *a* = 0.062 *mV*^−1^, *b* = 3.57, *g*_*SFA*_ = 100 *nS*, *V*_*K*_ = −80 *mV*, *τ*_*w*_ = 80 *ms, b*_*SFA*_ = 0.1.

### Synaptic Gating Variables

The synaptic gating variable 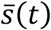 represents the fraction of open channels. For recurrent connections:

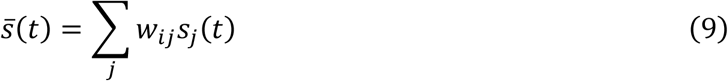

where *w*_*ij*_ is the synaptic weight from presynaptic neuron *j* to postsynaptic neuron *i*, and *s*_*j*_ (*t*) is the gating variable driven by presynaptic spikes.

For external inputs, 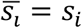, determined independently by each neuron’s input pulse sequence. For AMPA and GABA receptors, *s*_*X*_(*t*) evolves as:

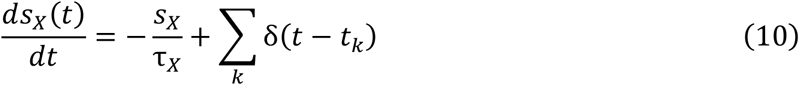

where *X* represents AMPA or GABA, and *t*_*k*_ denotes the presynaptic spike times.

NMDA receptor kinetics follow a two-stage process:

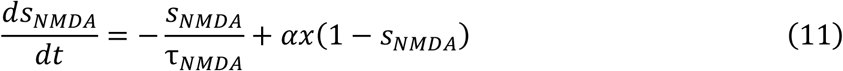

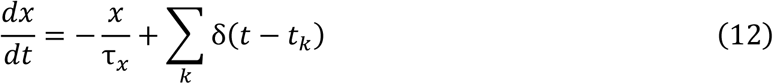

Here, *τ*_*AMPA*_ = 2 *ms, τ*_*GABA*_ = 5 *ms, τ*_*NMDA*_ = 100 *ms, τ*_*x*_ = 2 *ms, α* = 0.5 *kHz*.

### Background Inputs and Connectivity

All neurons receive independent Poisson background inputs via AMPA receptors, with firing rate *v*_*background*_.

The maximal conductances for background noises targeting excitatory and inhibitory neurons are *g*^*background*→*E*^ and *g*^*background*→*I*^, respectively.

Recurrent connection strength between two excitatory neurons depends on the difference between their preferred angles θ_*i*_ and θ_*j*_ :

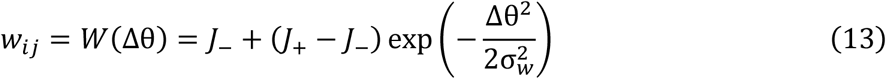

where *J*_+_ represents the peak synaptic strength, *J*_−_ is the baseline connectivity, and σ_*w*_ determines the spatial width of the connectivity profile. Here, *v*_*background*_ = 100 *Hz, g*^*background*→*E*^ = 1 *nS, g*^*background*→*I*^ = 1.62 *nS, J*_+_ = 35, *J*_−_ = 0, σ_*w*_ = 2.4.

### Cortical Decision-Making Module

The cortical decision-making module was implemented following the cortical network model proposed by Wang ^19^. This module was included to examine the benefits of combining two degenerate decision-making modules. It integrates evidence derived either from Purkinje neurons (in the cortico-cerebello-cortical model) or from the cortical continuous attractor network (in the cortico-cortical model).

The network consists of *N*_*E*_ excitatory pyramidal neurons and *N*_*I*_ inhibitory interneurons. Neuronal and synaptic dynamics follow the continuous attractor model described previously, with the following modifications to implement decision-making functionality.

The excitatory population is divided into three distinct subpopulations:

a. Non-selective population (*N*_0_) — responds only to background noise.
b. Selective population 1 (*N*_1_) — encodes the left decision, receiving input from Purkinje neurons or attractor neurons representing left-side cues.
c. Selective population 2 (*N*_2_) — encodes the right decision, receiving input from Purkinje neurons or attractor neurons representing right-side cues.

The size of the selective populations is determined by the fraction of selective neurons, *f*_*sel*_, such that *N*_1_ = *N*_2_ = *f*_*sel*_ *N*_*E*_.

All neurons receive global inhibition from the inhibitory pool, while excitatory connectivity is structured to promote competition. Synaptic weights between excitatory neurons, *w*_*ij*_, depend on population identity:

- Within the same selective population (recurrent excitation), *w*_+_ = *w*_*p*_.
- Between different selective populations, *w*_−_ = 1 − *f*_*sel*_ *w*_*p*_) /(1 − *f*_*sel*_).
- All other excitatory and inhibitory connections are set to unity (*w* = 1).

Each neuron receives independent Poisson background noise via AMPA receptors at a rate of *v*_*ext*_. Evidence signals originating from the cerebellum or the cortical continuous attractor module are modeled as external AMPA currents delivered to the corresponding selective populations. Here, *N*_*E*_ = 1600, *N*_*I*_ = 400, *f*_*sel*_ = 0.15, *w*_*p*_ = 1.7, *v*_*ext*_ = 2.4 k*Hz*.

### Cerebellar Model

The cerebellar circuit was modeled largely as a feedforward architecture. The network consists of three layers: (1) an input layer of mossy fibers, (2) a granule cell layer containing granule and Golgi cells, and (3) an output layer composed of Purkinje neurons.

Except in Figure 2, where each sensory stimulus is represented by a burst of spikes, the cerebellar model receives input from excitatory populations of the cortical continuous attractor module. The axon terminals of cortical neurons are modeled as mossy fibers, with a total of 60 fibers.

The granule cell layer contains *N*_*gc*_ excitatory granule cells and *N*_*Golgi*_ inhibitory Golgi cells. Granule cell membrane dynamics follow the adaptive quadratic integrate-and-fire (AdQIF) model:

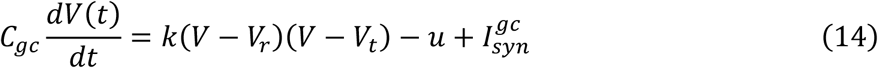

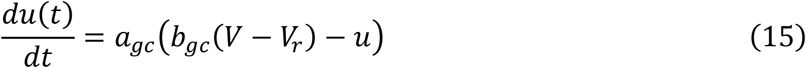

where *V* is the membrane potential and *u* the adaptation variable. *C*_*gc*_ is the membrane capacitance, k a scaling factor, *V*_*r*_ the resting potential, and *V*_*t*_ the threshold parameter. Adaptation dynamics are controlled by the time constant *a*_*gc*_ and sensitivity parameter *b*_*gc*_. When *V* reaches *V*_*peak*_, the potential is reset to *c*_*reset*_ and *u* is incremented by *d*_*reset*_.

The total synaptic current is:

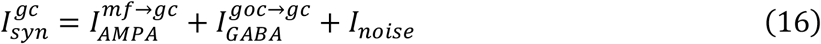

AMPA, GABA, and background Poisson noise currents are computed as in the cortical model.Golgi cells provide feedback inhibition to regulate granule cell gain and temporal dynamics. They are modeled as leaky integrate‐and‐fire (LIF) neurons:

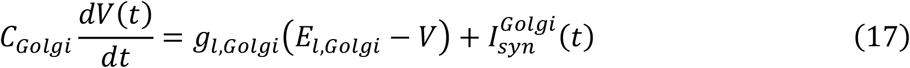

where *C*_*Golgi*_ is the membrane capacitance and *g*_*l,Golgi*_ is the leak conductance. The total synaptic current is:

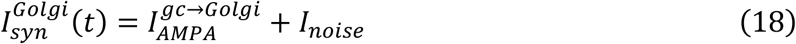

The output layer comprises *N*_*PN*_ Purkinje neurons (half per side), modeled using the adaptive exponential integrate‐and‐fire (AdEx) formalism with spike‐triggered adaptation and colored noise ^43^:

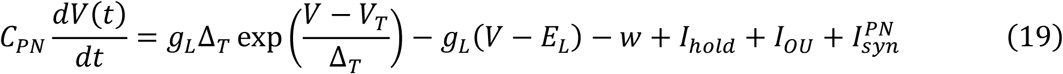

where *C*_*PN*_ is the membrane capacitance, *g*_*L*_ the leak conductance, and *E*_*L*_ the leak reversal potential. The exponential term models the rapid depolarization during spike generation, with slope factor Δ_*T*_ and threshold *V*_*T*_.

The adaptation variable *w* follows:

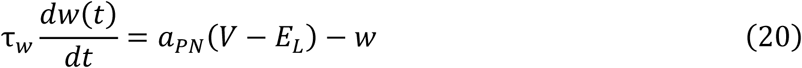

When *V* reaches 60 mV, a spike is emitted, *V* is reset to *V*_*reset*_, and *w* is incremented by *b*_*PN*_. Intrinsic background activity is modeled using Ornstein-Uhlenbeck (OU) noise:

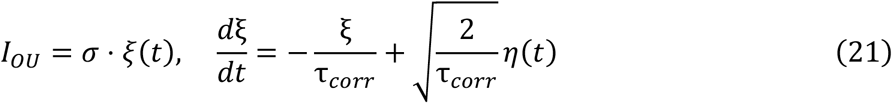

where *σ* is the noise standard deviation, *τ*_*corr*_ the correlation time constant, and *η*(*t*) Gaussian white noise. A constant holding current *I*_*hold*_ modulates baseline excitability. For simplicity, cerebellar nuclei neurons are not modeled explicitly; their function is merged into the Purkinje neuron population, which acts as an excitatory output to downstream cortical circuits.

Each granule cell receives excitatory input from four randomly selected mossy fibers. Granule cell and Golgi cells form a feedback inhibition loop:

- granule → Golgi connections (AMPA): *p*_*gc*→*Golgi*_ = 0.1
- Golgi → granule connections (GABA): *p*_*Golgi*→*gc*_ = 0.5

This loop regulates granule cell gain and temporal sparsity. Granule cells project to Purkinje neurons via parallel fibers. The synaptic weights *w*_*PF*−*PN*_ are plastic, following a spike‐timing‐dependent plasticity (STDP) rule:

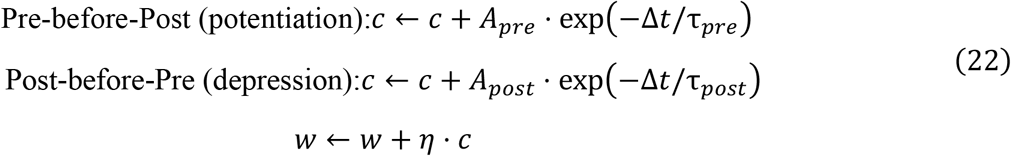

where η is the learning rate, and *τ*_*pre*_ and *τ*_*post*_ define the temporal windows for plasticity. *c* denotes the synaptic eligibility trace. Here, *C*_*gc*_ = 100 *pF*, *V*_*r*_ = −60 *mV*, *V*_*t*_ = −40 *mV*, k = 0.7 *pA*/(*mV*^2^), *a*_*gc*_ = 0.03 *ms*^−1^, *b*_*gc*_ = −2 *nS*, *C*_*Golgi*_ = 50 *pF*, *g*_*l,Golgi*_ = 5 *nS*, *E*_*l,Golgi*_ = −65 *mV*, *N*_*PC*_ = 400, *C*_*PC*_ = 268 *pF*, *g*_*L*_ = 8.47 *nS*, Δ_*T*_ = 0.85 *mV*, *V*_*T*_ = −53.23 *mV*, *E*_*L*_ = −51.31 *mV*, *τ*_*w*_ = 20.76 *ms*, *a*_*PC*_ = 37.79 *nS*, *b*_*PC*_ = 441.12 *pA*, *V*_*reset*_ = −60.35 *mV*, σ = 40 *pA, τ*_*corr*_ = 2 *ms, I*_*hold*_ = −140 *pA*.

### Sensitivity to Mutual Inhibition between Purkinje Neurons

Mutual inhibition between Purkinje neurons is critical for evidence competition. Excessively high inhibition conductance destabilizes baseline firing, producing spontaneous bistability, which is biologically unrealistic ^45^. A conductance value is considered biologically plausible if the firing‐rate difference between the two Purkinje neuron populations remains below 2 Hz after 6500 ms of simulation without external stimulation.

The fraction of completed trials was defined as the ratio of valid trials to total trials. To handle undecided trials conservatively, we followed the standard practice of random assignment to left or right decisions while also reporting the fraction of completed trials as an independent metric.

### t‐SNE Visualization of Granule Cell Population Activity

To visualize high‐dimensional granule cell activity across sparsity levels, we applied t‐Distributed Stochastic Neighbor Embedding (t‐SNE). Spike trains from N = 2000 granule cells were converted into population state vectors using non‐overlapping 20 ms bins. Each vector contained spike counts across all recorded neurons. State vectors from left‐ and right‐dominant conditions were merged for joint analysis.

t‐SNE was implemented using Scikit‐learn with the following parameters: Components = 3; Perplexity = 15; Initialization = PCA; Distance metric = Euclidean

This configuration balances local/global structure preservation and ensures stable manifold visualization.

### Population Firing Rate

Instantaneous population firing rates were computed using a 100 ms sliding window moving in 2 ms steps. Within each window, total spikes were divided by the number of neurons and the window duration to obtain population firing rates.

### Simulations

All simulations were performed on a Linux workstation. The coupled differential equations describing neuronal and synaptic dynamics were numerically integrated using a modified second‐order Runge–Kutta (RK2) algorithm with a time step of 0.02 ms.

Decision‐making performance was evaluated as the average percentage of correct choices across n = 200 trials for each parameter set and left-right stimulus condition.

## Acknowledgements

This work is supported by the National Key Research and Development Program of China (2023YFF1204200) and the National Natural Science Foundation of China (62476197, 12372060).

## Supplementary Figures

**Figure S1.**
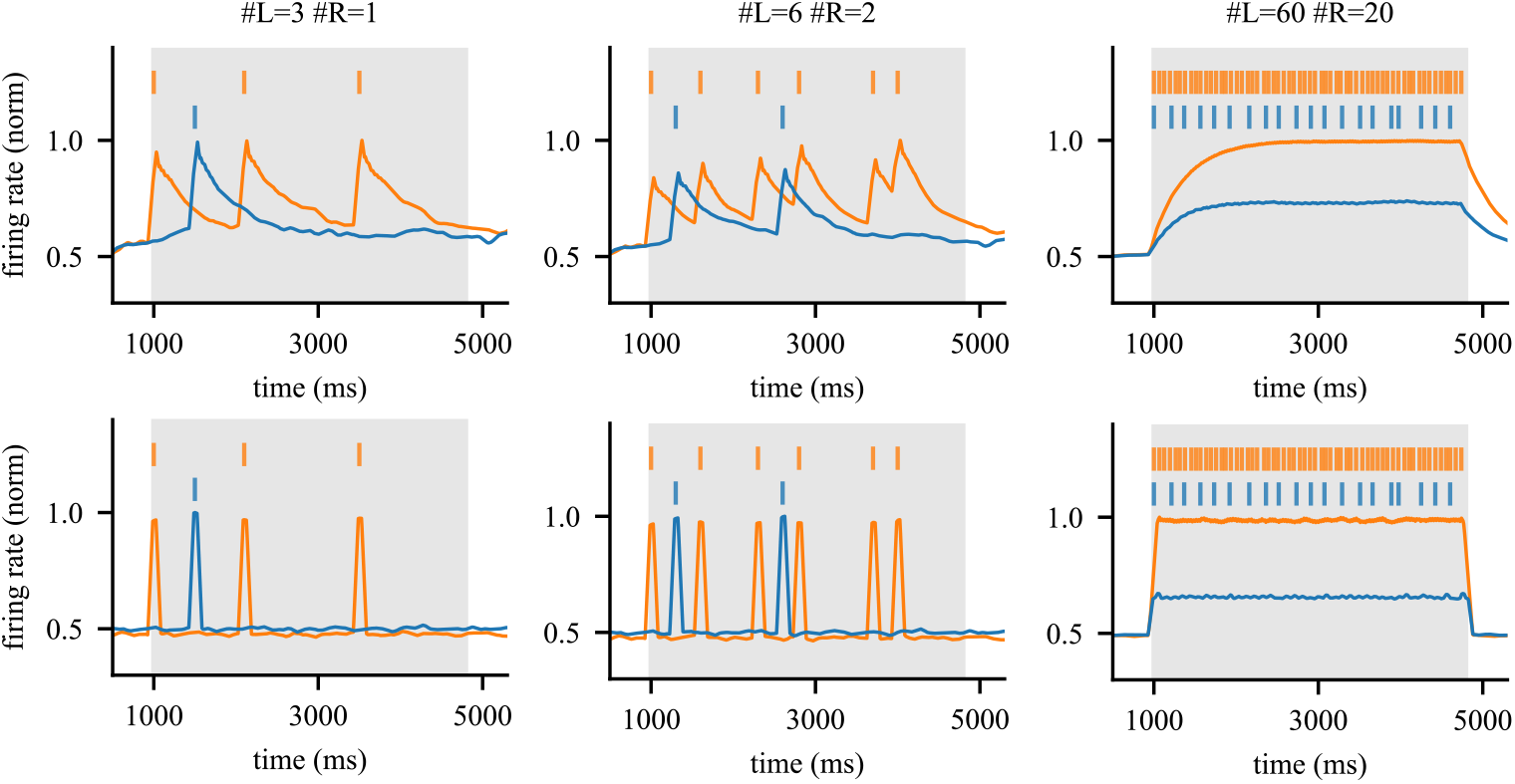
Purkinje neuron firing rate accumulation dynamics (type II, top; type I, bottom) without mutual inhibition across increasing stimulus frequencies. Colored vertical lines mark individual stimulus, while colored traces show the corresponding firing‐rate responses of Purkinje neurons on each side.

**Figure S2.**
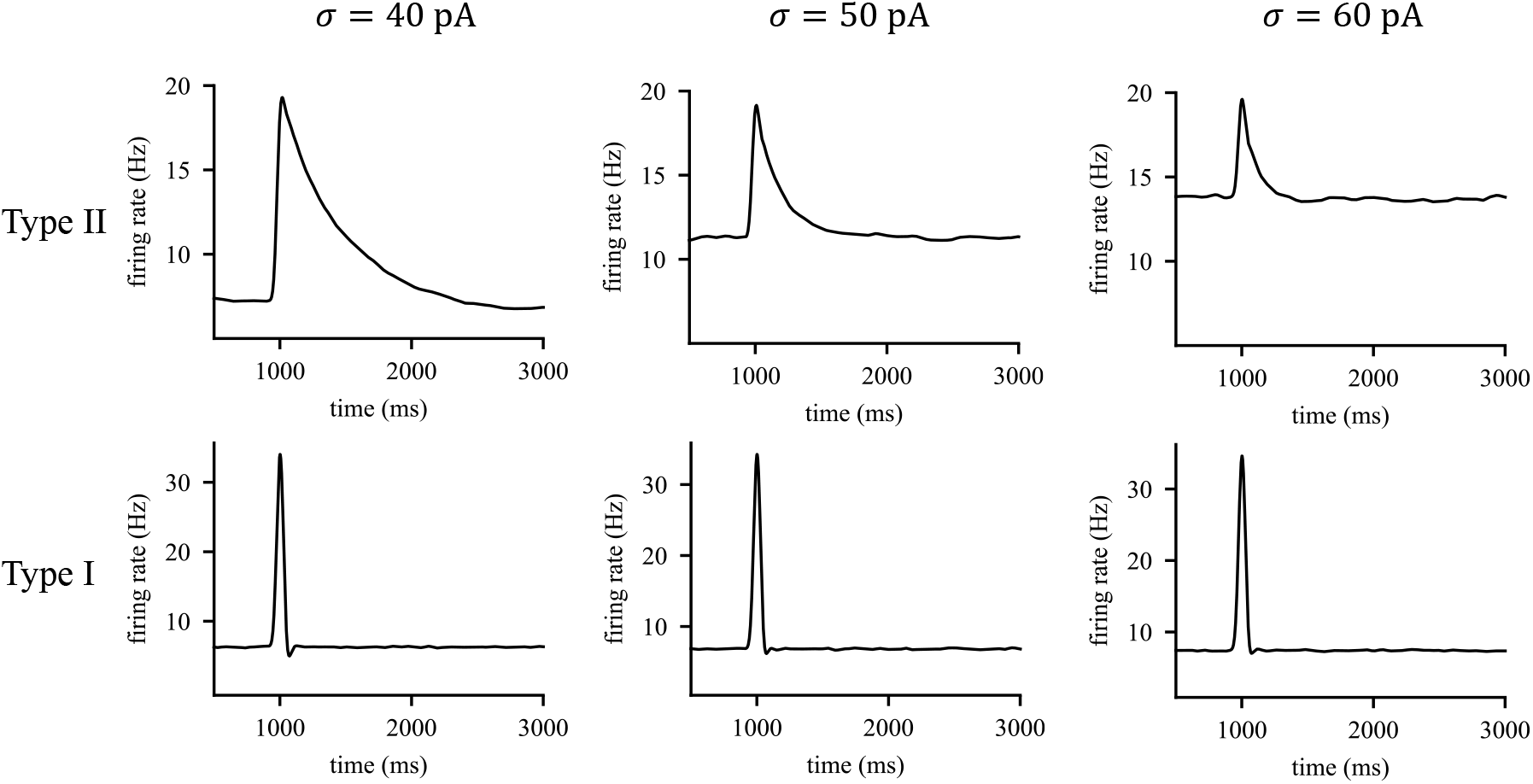
Example population firing rates of Purkinje neurons at different noise variances. Population firing‐rate responses are shown for the Type II model (top) and Type I model (bottom) under varying levels of input noise. I_hold_ = ‐140 pA.

**Figure S3.**
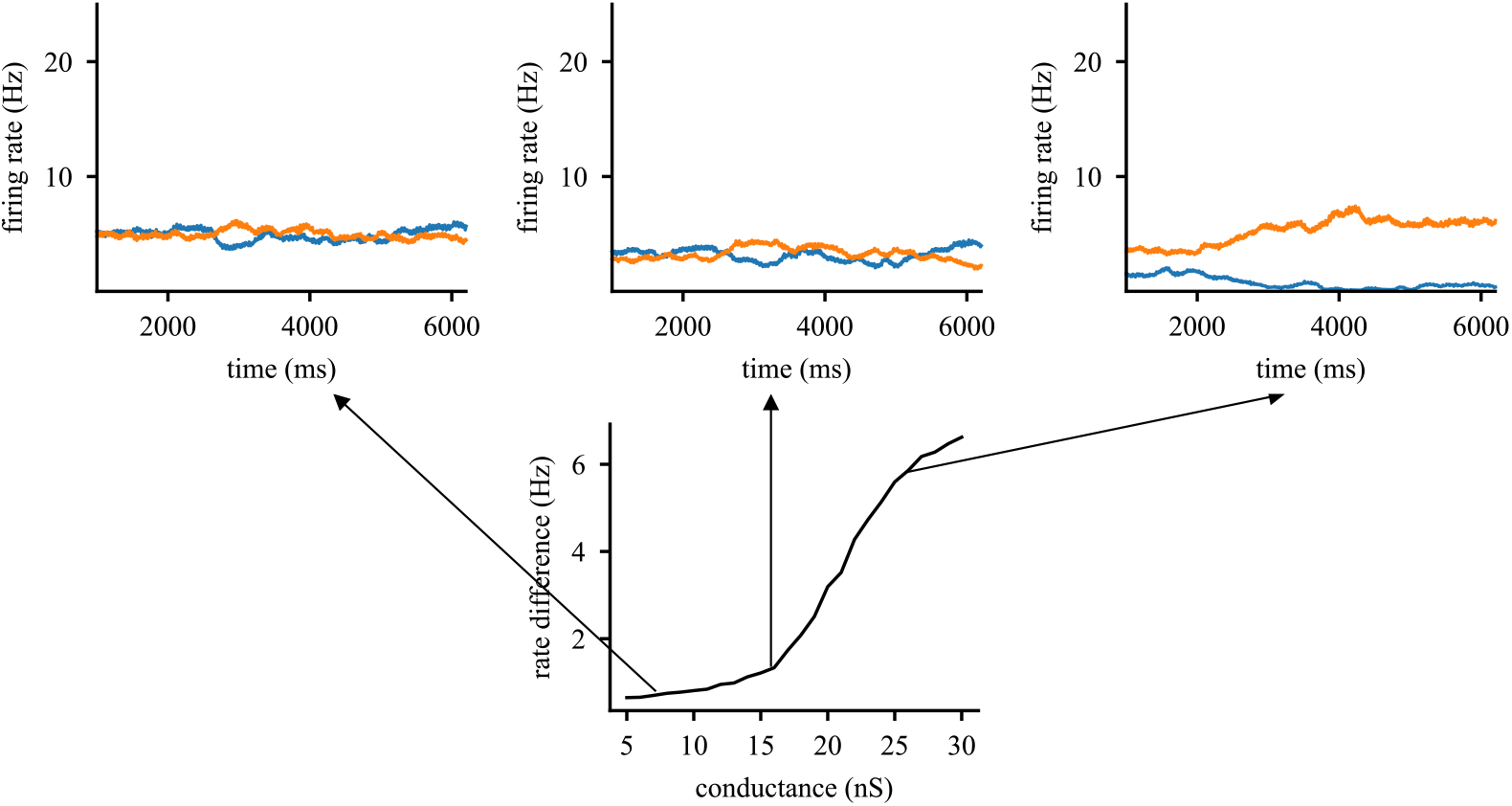
Instability of Purkinje neuron spontaneous firing dynamics at large mutual inhibition strengths. Increasing mutual inhibition beyond a critical threshold leads to unstable spontaneous firing behaviors in Purkinje neurons.

**Figure S4.**
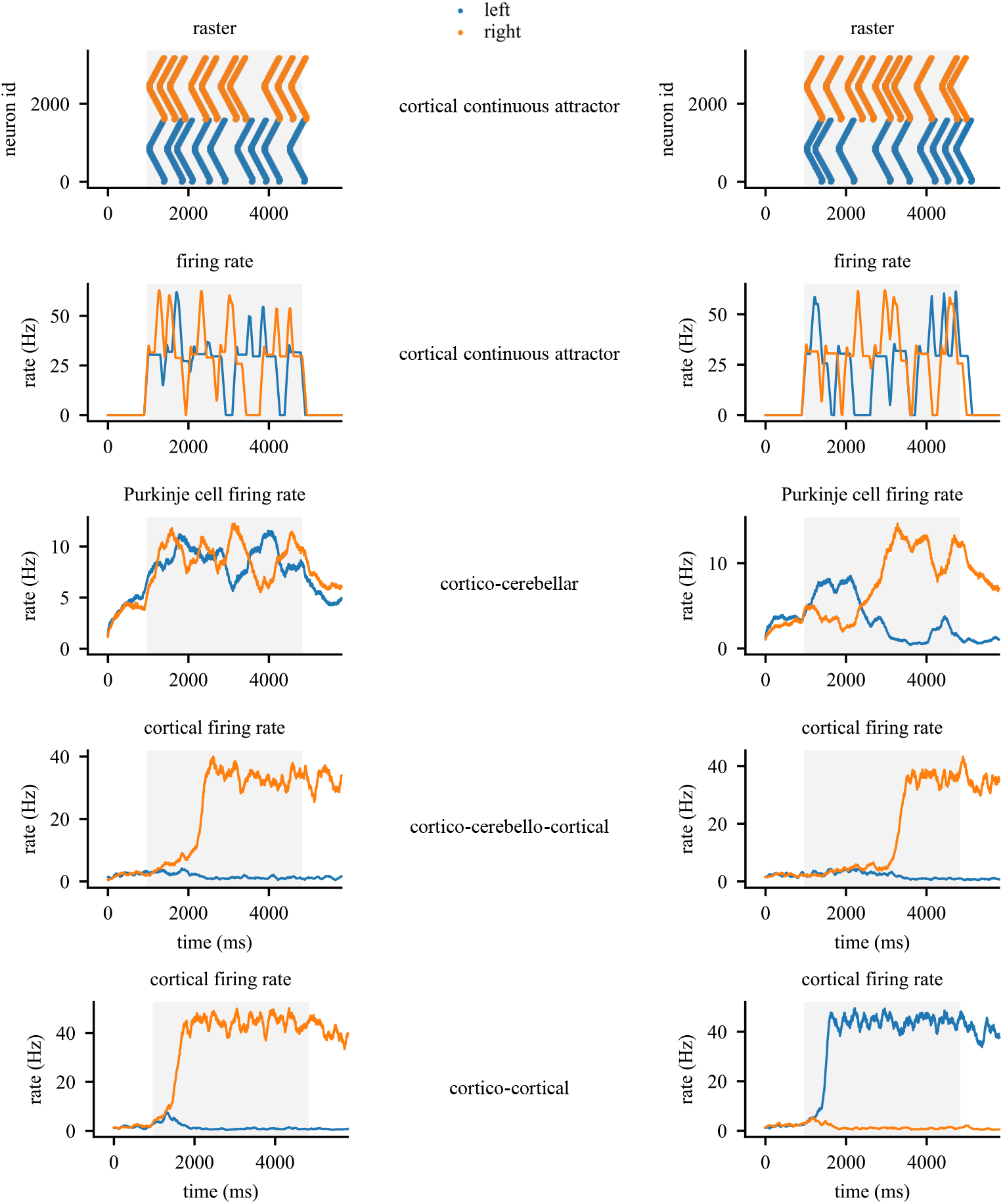
Example circuit dynamics illustrating complementary roles of cerebellar and cortical decision modules. From top to bottom: spike raster in the cortical continuous-attractor module, corresponding population firing rates, Purkinje-neuron population firing rates in the cortico–cerebellar model, cortical population firing rates in the cortico–cerebello–cortical model, and cortical population firing rates in the cortico– cortical model. In both columns, evidence favors the left choice. In the left example, the cortico– cerebellar model fails to reach a categorical decision, whereas both the cortico–cortical and cortico– cerebello–cortical models choose correctly. In the right example, the cortico–cortical model chooses incorrectly due to a primacy bias, whereas both the cortico–cerebellar and cortico–cerebello–cortical models choose correctly.

**Figure S5.**
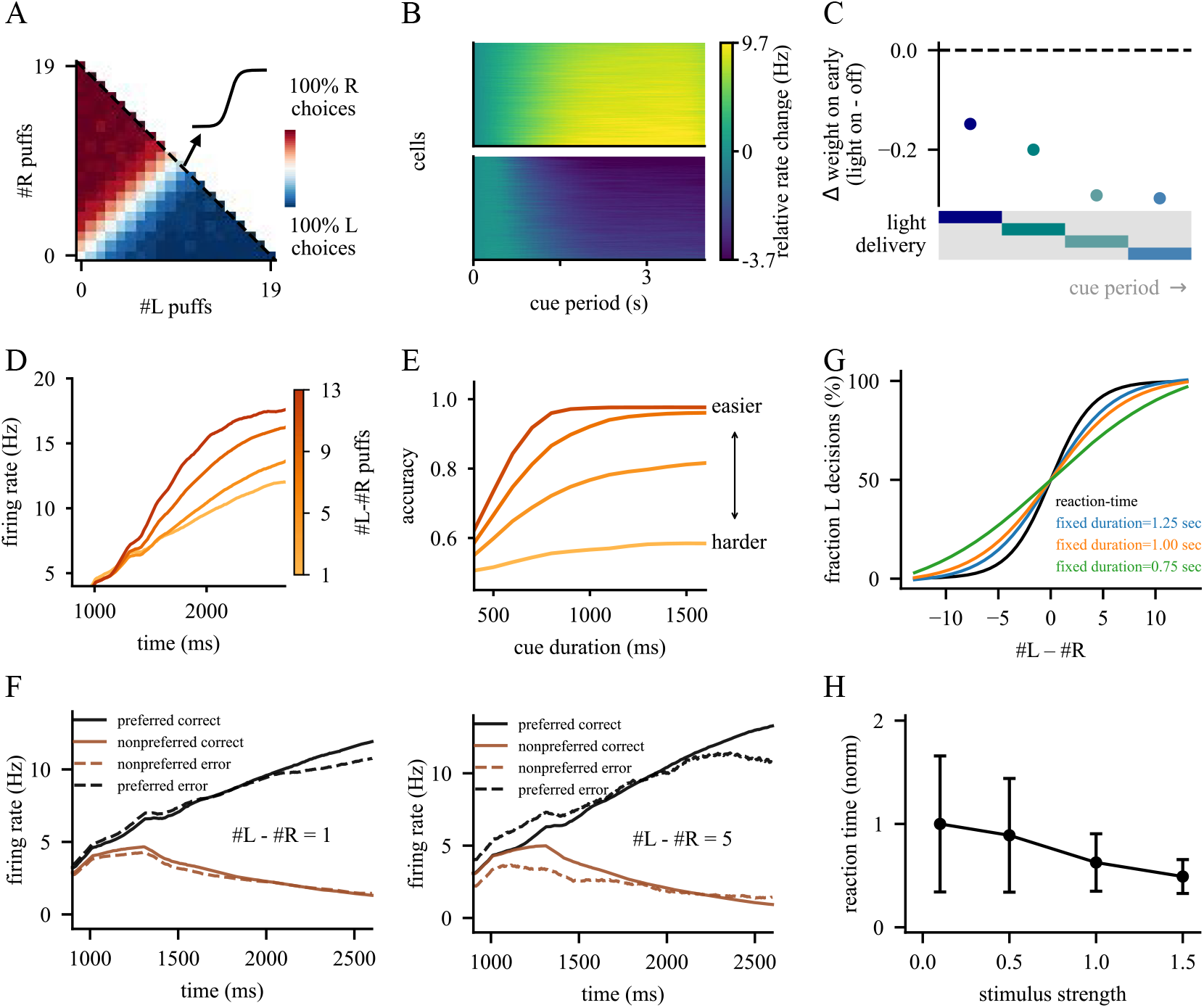
Evidence-accumulation dynamics can also be captured by the cortico–cerebellar model. (A) Choice probability as a function of left and right puff counts. (B) Cue-period activity of winning (top) and losing (bottom) Purkinje neuron populations. (C) Weight change following cue-period Purkinje neuron perturbations applied at different times. (D) Winning-population Purkinje neuron trajectories across evidence differences. (E) Psychometric curves across cue durations and difficulty levels. (F) Population trajectories for correct and error trials across stimulus differences. (G) Psychometric curves comparing different fixed cue durations with the reaction-time paradigm. (H) Mean reaction time changes versus stimulus strength. Panels A–F: fixed-time; panel H: reaction-time; panel G: both. Compared with Figure 7, panels B and D are identical; all other panels differ.

**Figure S6.**
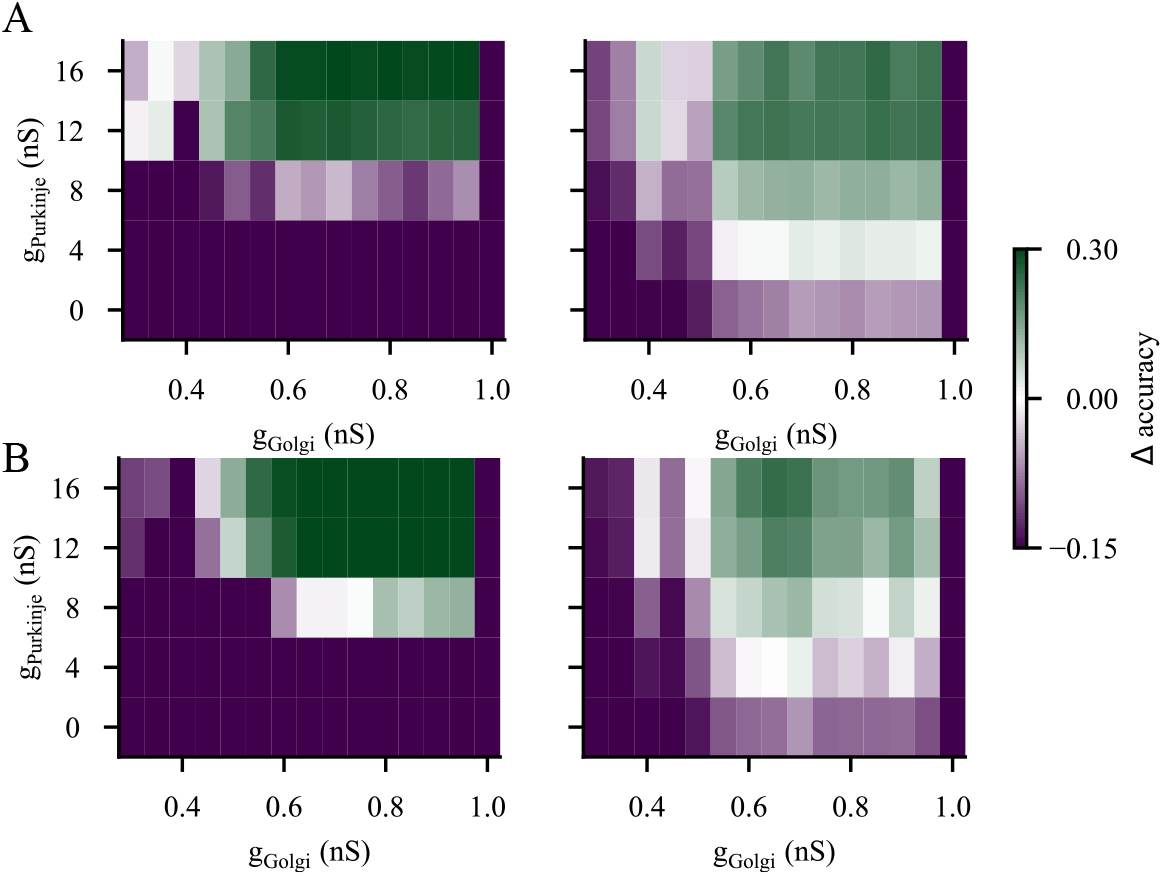
Accuracy‐gain heatmaps as a function of Golgi cell and Purkinje neuron inhibition strengths at overlap ratios of 40% (A) and 60% (B) Heatmaps show the relative improvement in decision accuracy for different inhibitory strengths. Left panels depict cortico‐cerebellar minus cortico‐cortical performance; right panels depict cortico‐cerebello‐cortical minus cortico‐cortical performance.

**Figure S7.**
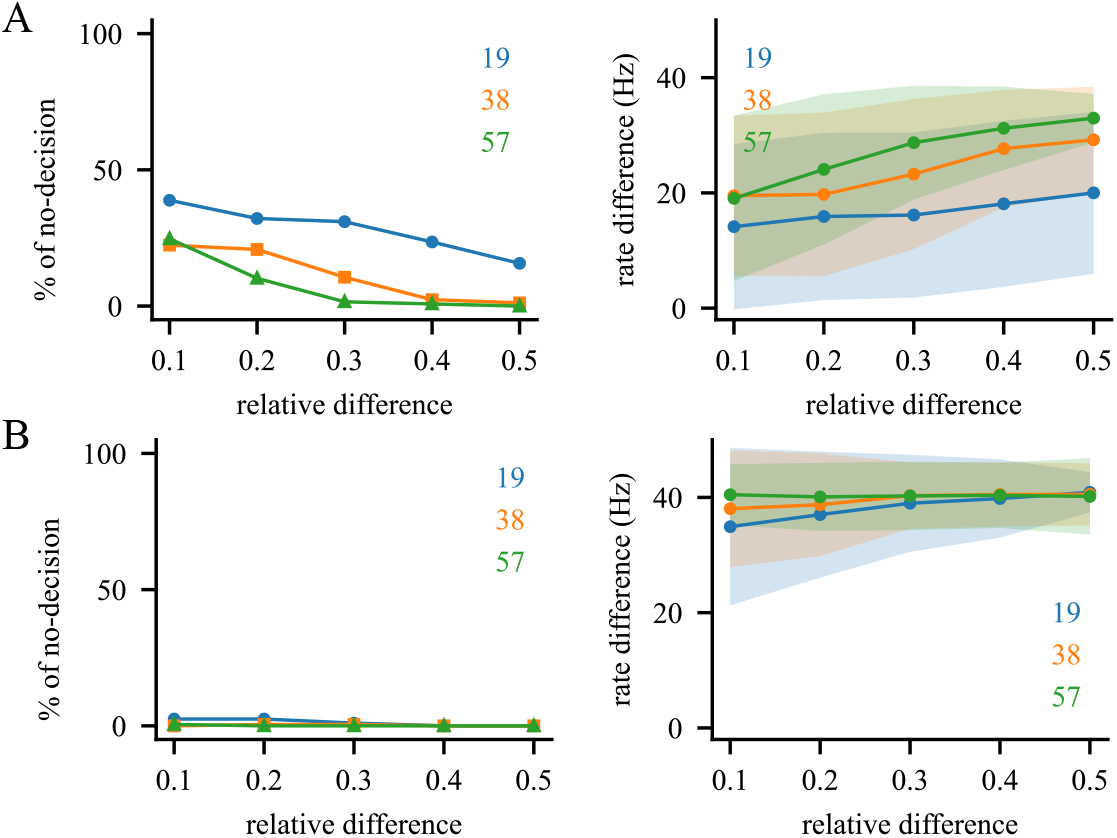
Cortical preprocessing enhances evidence accumulation in the cortico‐cerebello‐cortical Model. (A) Proportion of undecided trials (left) and firing‐rate difference between opposing cortical populations at the end of the cue period in response to direct sensory‐triggered spike bursts. (B) Same analysis as in (A), but for cortical‐preprocessed inputs, showing improved decision formation and reduced indecision. Colored numbers indicate total evidence across both sides.

## Notes

### Competing Interest Statement

The authors have declared no competing interest.

## References

1. Marr, D. (1969). A theory of cerebellar cortex. J Physiol 202, 437–470. 10.1113/jphysiol.1969.sp008820.

2. Albus, J.S. (1971). A theory of cerebellar function. Math Biosci 10, 25–61. 10.1016/0025-5564(71)90051-4.

3. Barri, A., Wiechert, M.T., Jazayeri, M., and DiGregorio, D.A. (2022). Synaptic basis of a sub-second representation of time in a neural circuit model. Nat Commun 13, 7902. 10.1038/s41467-022-35395-y.

4. Bhasin, B.J., Raymond, J.L., and Goldman, M.S. (2024). Synaptic weight dynamics underlying memory consolidation: Implications for learning rules, circuit organization, and circuit function. Proc Natl Acad Sci U S A 121, e2406010121. 10.1073/pnas.2406010121.

5. Fakharian, M.A., Shoup, A.M., Hage, P., Elseweifi, H.Y., and Shadmehr, R. (2025). A vector calculus for neural computation in the cerebellum. Science 388, 869–875. 10.1126/science.adu6331.

6. Suvrathan, A., Payne, H.L., and Raymond, J.L. (2016). Timing Rules for Synaptic Plasticity Matched to Behavioral Function. Neuron 92, 959–967. 10.1016/j.neuron.2016.10.022.

7. International Brain, L., Angelaki, D., Benson, B., Benson, J., Birman, D., Bonacchi, N., Bougrova, K., Bruijns, S.A., Carandini, M., Catarino, J.A., et al. (2025). A brain-wide map of neural activity during complex behaviour. Nature 645, 177–191. 10.1038/s41586-025-09235-0.

8. Wagner, M.J., and Luo, L. (2020). Neocortex-Cerebellum Circuits for Cognitive Processing. Trends Neurosci 43, 42–54. 10.1016/j.tins.2019.11.002.

9. Deverett, B., Koay, S.A., Oostland, M., and Wang, S.S. (2018). Cerebellar involvement in an evidence-accumulation decision-making task. Elife 7, e36781. 10.7554/eLife.36781.

10. Deverett, B., Kislin, M., Tank, D.W., and Wang, S.S. (2019). Cerebellar disruption impairs working memory during evidence accumulation. Nat Commun 10, 3128. 10.1038/s41467-019-11050-x.

11. Carta, I., Chen, C.H., Schott, A.L., Dorizan, S., and Khodakhah, K. (2019). Cerebellar modulation of the reward circuitry and social behavior. Science 363. 10.1126/science.aav0581.

12. Diedrichsen, J., King, M., Hernandez-Castillo, C., Sereno, M., and Ivry, R.B. (2019). Universal Transform or Multiple Functionality? Understanding the Contribution of the Human Cerebellum across Task Domains. Neuron 102, 918–928. 10.1016/j.neuron.2019.04.021.

13. Zang, Y., and De Schutter, E. (2023). Recent data on the cerebellum require new models and theories. Curr Opin Neurobiol 82, 102765. 10.1016/j.conb.2023.102765.

14. Zhang, X.Y., Wu, W.X., Shen, L.P., Ji, M.J., Zhao, P.F., Yu, L., Yin, J., Xie, S.T., Xie, Y.Y., Zhang, Y.X., et al. (2024). A role for the cerebellum in motor-triggered alleviation of anxiety. Neuron 112, 1165–1181 e1168. 10.1016/j.neuron.2024.01.007.

15. Koay, S.A., Charles, A.S., Thiberge, S.Y., Brody, C.D., and Tank, D.W. (2022). Sequential and efficient neural-population coding of complex task information. Neuron 110, 328–349 e311. 10.1016/j.neuron.2021.10.020.

16. Harvey, C.D., Coen, P., and Tank, D.W. (2012). Choice-specific sequences in parietal cortex during a virtual-navigation decision task. Nature 484, 62–68. 10.1038/nature10918.

17. Nieh, E.H., Schottdorf, M., Freeman, N.W., Low, R.J., Lewallen, S., Koay, S.A., Pinto, L., Gauthier, J.L., Brody, C.D., and Tank, D.W. (2021). Geometry of abstract learned knowledge in the hippocampus. Nature 595, 80–84. 10.1038/s41586-021-03652-7.

18. Brunton, B.W., Botvinick, M.M., and Brody, C.D. (2013). Rats and humans can optimally accumulate evidence for decision-making. Science 340, 95–98. 10.1126/science.1233912.

19. Wang, X.J. (2002). Probabilistic decision making by slow reverberation in cortical circuits. Neuron 36, 955–968. 10.1016/s0896-6273(02)01092-9.

20. Goldman, M.S. (2017). Memory without Feedback in a Neural Network. Neuron 93, 715. 10.1016/j.neuron.2017.01.007.

21. Machens, C.K., Romo, R., and Brody, C.D. (2005). Flexible control of mutual inhibition: a neural model of two-interval discrimination. Science 307, 1121–1124. 10.1126/science.1104171.

22. Bi, Z., and Zhou, C. (2020). Understanding the computation of time using neural network models. Proc Natl Acad Sci U S A 117, 10530–10540. 10.1073/pnas.1921609117.

23. Nguyen, T.M., Thomas, L.A., Rhoades, J.L., Ricchi, I., Yuan, X.C., Sheridan, A., Hildebrand, D.G.C., Funke, J., Regehr, W.G., and Lee, W.A. (2023). Structured cerebellar connectivity supports resilient pattern separation. Nature 613, 543–549. 10.1038/s41586-022-05471-w.

24. Roth, A., and Häusser, M. (2001). Compartmental models of rat cerebellar Purkinje cells based on simultaneous somatic and dendritic patch-clamp recordings. J Physiol 535, 445–472. 10.1111/j.1469-7793.2001.00445.x.

25. Ohtsuki, G., Piochon, C., Adelman, J.P., and Hansel, C. (2012). SK2 channel modulation contributes to compartment-specific dendritic plasticity in cerebellar Purkinje cells. Neuron 75, 108–120. 10.1016/j.neuron.2012.05.025.

26. Cayco-Gajic, N.A., Clopath, C., and Silver, R.A. (2017). Sparse synaptic connectivity is required for decorrelation and pattern separation in feedforward networks. Nat Commun 8, 1116. 10.1038/s41467-017-01109-y.

27. Babadi, B., and Sompolinsky, H. (2014). Sparseness and expansion in sensory representations. Neuron 83, 1213–1226. 10.1016/j.neuron.2014.07.035.

28. Litwin-Kumar, A., Harris, K.D., Axel, R., Sompolinsky, H., and Abbott, L.F. (2017). Optimal Degrees of Synaptic Connectivity. Neuron 93, 1153–1164 e1157. 10.1016/j.neuron.2017.01.030.

29. Pham, T., and Hansel, C. (2023). Intrinsic threshold plasticity: cholinergic activation and role in the neuronal recognition of incomplete input patterns. J Physiol 601, 3221–3239. 10.1113/JP283473.

30. Rancz, E.A., Ishikawa, T., Duguid, I., Chadderton, P., Mahon, S., and Häusser, M. (2007). High-fidelity transmission of sensory information by single cerebellar mossy fibre boutons. Nature 450, 1245–1248. 10.1038/nature05995.

31. Watt, A.J., Cuntz, H., Mori, M., Nusser, Z., Sjostrom, P.J., and Häusser, M. (2009). Traveling waves in developing cerebellar cortex mediated by asymmetrical Purkinje cell connectivity. Nat Neurosci 12, 463–473. 10.1038/nn.2285.

32. Witter, L., Rudolph, S., Pressler, R.T., Lahlaf, S.I., and Regehr, W.G. (2016). Purkinje Cell Collaterals Enable Output Signals from the Cerebellar Cortex to Feed Back to Purkinje Cells and Interneurons. Neuron 91, 312–319. 10.1016/j.neuron.2016.05.037.

33. Zang, Y., Hong, S., and De Schutter, E. (2020). Firing rate-dependent phase responses of Purkinje cells support transient oscillations. Elife 9, e60692. 10.7554/eLife.60692.

34. Roš, H., Xie, Y., Sadeh, S., and Silver, R.A. (2025). Population activity of mossy fibre axon input to the cerebellar cortex during behaviours. bioRxiv, 2025.2003.2023.644738. 10.1101/2025.03.23.644738.

35. Pagan, M., Tang, V.D., Aoi, M.C., Pillow, J.W., Mante, V., Sussillo, D., and Brody, C.D. (2025). Individual variability of neural computations underlying flexible decisions. Nature 639, 421–429. 10.1038/s41586-024-08433-6.

36. Schall, J.D. (2001). Neural basis of deciding, choosing and acting. Nat Rev Neurosci 2, 33–42. 10.1038/35049054.

37. Bender, F., Sermet, B.S., Borda Bossana, S., Barri, A., Schamiloglu, S., Diana, G., Costreie, M.-M., Moneron, G., Hantman, A.W., and DiGregorio, D.A. (2026). Synaptic dynamics as a tunable substrate shaping neuronal activity sequences. bioRxiv, 2026.2003.2022.713064. 10.64898/2026.03.22.713064.

38. Wagner, M.J., Kim, T.H., Kadmon, J., Nguyen, N.D., Ganguli, S., Schnitzer, M.J., and Luo, L. (2019). Shared Cortex-Cerebellum Dynamics in the Execution and Learning of a Motor Task. Cell 177, 669–682 e624. 10.1016/j.cell.2019.02.019.

39. Gao, Z., Davis, C., Thomas, A.M., Economo, M.N., Abrego, A.M., Svoboda, K., De Zeeuw, C.I., and Li, N. (2018). A cortico-cerebellar loop for motor planning. Nature 563, 113–116. 10.1038/s41586-018-0633-x.

40. Ren, Z., Wang, X., Angelov, M., De Zeeuw, C.I., and Gao, Z. (2025). Neuronal dynamics of cerebellum and medial prefrontal cortex in adaptive motor timing. Nat Commun 16, 612. 10.1038/s41467-025-55884-0.

41. Zang, Y., and De Schutter, E. (2021). The Cellular Electrophysiological Properties Underlying Multiplexed Coding in Purkinje Cells. J Neurosci 41, 1850–1863. 10.1523/JNEUROSCI.1719-20.2020.

42. Zang, Y., Dieudonne, S., and De Schutter, E. (2018). Voltage- and Branch-Specific Climbing Fiber Responses in Purkinje Cells. Cell Rep 24, 1536–1549. 10.1016/j.celrep.2018.07.011.

43. Buchin, A., Rieubland, S., Häusser, M., Gutkin, B.S., and Roth, A. (2016). Inverse Stochastic Resonance in Cerebellar Purkinje Cells. PLoS Comput Biol 12, e1005000. 10.1371/journal.pcbi.1005000.

44. Nolan, M.F., Malleret, G., Lee, K.H., Gibbs, E., Dudman, J.T., Santoro, B., Yin, D., Thompson, R.F., Siegelbaum, S.A., Kandel, E.R., and Morozov, A. (2003). The Hyperpolarization-Activated HCN1 Channel Is Important for Motor Learning and Neuronal Integration by Cerebellar Purkinje Cells. Cell 115, 551–564. 10.1016/S0092-8674(03)00884-5.

45. Schonewille, M., Khosrovani, S., Winkelman, B.H.J., Hoebeek, F.E., De Jeu, M.T.G., Larsen, I.M., Van Der Burg, J., Schmolesky, M.T., Frens, M.A., and De Zeeuw, C.I. (2006). Purkinje cells in awake behaving animals operate at the upstate membrane potential. Nat Neurosci 9, 459–461. 10.1038/nn0406-459.

46. Huk, A.C., and Shadlen, M.N. (2005). Neural activity in macaque parietal cortex reflects temporal integration of visual motion signals during perceptual decision making. J Neurosci 25, 10420–10436. 10.1523/JNEUROSCI.4684-04.2005.

47. Kiani, R., Hanks, T.D., and Shadlen, M.N. (2008). Bounded Integration in Parietal Cortex Underlies Decisions Even When Viewing Duration Is Dictated by the Environment. J Neurosci 28, 3017–3029. 10.1523/jneurosci.4761-07.2008.

48. Lam, N.H., Borduqui, T., Hallak, J., Roque, A., Anticevic, A., Krystal, J.H., Wang, X.J., and Murray, J.D. (2022). Effects of Altered Excitation-Inhibition Balance on Decision Making in a Cortical Circuit Model. J Neurosci 42, 1035–1053. 10.1523/JNEUROSCI.1371-20.2021.

49. Gupta, D., Kopec, C.D., Bondy, A.G., Luo, T.Z., Elliott, V., and Brody, C.D. (2026). A multi-region recurrent circuit for evidence accumulation in rats. Neuron 114, 521–535 e525. 10.1016/j.neuron.2025.12.029.

50. Brody, C.D., and Hanks, T.D. (2016). Neural underpinnings of the evidence accumulator. Curr Opin Neurobiol 37, 149–157. 10.1016/j.conb.2016.01.003.

51. Roitman, J.D., and Shadlen, M.N. (2002). Response of Neurons in the Lateral Intraparietal Area during a Combined Visual Discrimination Reaction Time Task. J Neurosci 22, 9475–9489. 10.1523/jneurosci.22-21-09475.2002.

52. Strick, P.L., Dum, R.P., and Fiez, J.A. (2009). Cerebellum and nonmotor function. Annu Rev Neurosci 32, 413–434. 10.1146/annurev.neuro.31.060407.125606.

53. Shadlen, M.N., and Newsome, W.T. (2001). Neural basis of a perceptual decision in the parietal cortex (area LIP) of the rhesus monkey. J Neurophysiol 86, 1916–1936. 10.1152/jn.2001.86.4.1916.

54. Lisberger, S.G., and Fuchs, A.F. (1978). Role of primate flocculus during rapid behavioral modification of vestibuloocular reflex. II. Mossy fiber firing patterns during horizontal head rotation and eye movement. J Neurophysiol 41, 764–777. 10.1152/jn.1978.41.3.764.

55. Ishikawa, T., Shimuta, M., and Häusser, M. (2015). Multimodal sensory integration in single cerebellar granule cells in vivo. Elife 4. 10.7554/eLife.12916.

56. Albantakis, L., and Deco, G. (2009). The encoding of alternatives in multiple-choice decision making. Proc Natl Acad Sci U S A 106, 10308–10313. 10.1073/pnas.0901621106.

57. Wong, K.F., Huk, A.C., Shadlen, M.N., and Wang, X.J. (2007). Neural circuit dynamics underlying accumulation of time-varying evidence during perceptual decision making. Front Comput Neurosci 1, 6. 10.3389/neuro.10.006.2007.

58. Ledergerber, D., and Larkum, M.E. (2010). Properties of layer 6 pyramidal neuron apical dendrites. J Neurosci 30, 13031–13044. 10.1523/JNEUROSCI.2254-10.2010.

59. Miller, P., Brody, C.D., Romo, R., and Wang, X.J. (2003). A recurrent network model of somatosensory parametric working memory in the prefrontal cortex. Cereb Cortex 13, 1208–1218. 10.1093/cercor/bhg101.

60. Spirou, G.A., Davis, K.A., Nelken, I., and Young, E.D. (1999). Spectral integration by type II interneurons in dorsal cochlear nucleus. J Neurophysiol 82, 648–663. 10.1152/jn.1999.82.2.648.

61. Beurrier, C., Bioulac, B., and Hammond, C. (2000). Slowly inactivating sodium current (I(NaP)) underlies single-spike activity in rat subthalamic neurons. J Neurophysiol 83, 1951–1957. 10.1152/jn.2000.83.4.1951.

62. de Solages, C., Szapiro, G., Brunel, N., Hakim, V., Isope, P., Buisseret, P., Rousseau, C., Barbour, B., and Lena, C. (2008). High-frequency organization and synchrony of activity in the purkinje cell layer of the cerebellum. Neuron 58, 775–788. 10.1016/j.neuron.2008.05.008.

63. Brown, L.S., Cho, J.R., Bolkan, S.S., Nieh, E.H., Schottdorf, M., Tank, D.W., Brody, C.D., Witten, I.B., and Goldman, M.S. (2026). Neural circuit models for evidence accumulation through choice-selective sequences. Nat Commun. 10.1038/s41467-026-70267-9.

64. Cayco-Gajic, N.A., and Silver, R.A. (2019). Re-evaluating Circuit Mechanisms Underlying Pattern Separation. Neuron 101, 584–602. 10.1016/j.neuron.2019.01.044.

65. Yu, L., Yang, Z., Bao, Y., and Zang, Y. (2026). Balancing Inhibition and Sparsity for Stable, Accurate Cerebellar Learning. bioRxiv, 2026.2004.2009.717611. 10.64898/2026.04.09.717611.

66. Thibaut, F. (2016). Basal ganglia play a crucial role in decision making. Dialogues Clin Neurosci 18, 3. 10.31887/DCNS.2016.18.1/fthibaut.

67. Mitchell, A.S. (2015). The mediodorsal thalamus as a higher order thalamic relay nucleus important for learning and decision-making. Neurosci Biobehav Rev 54, 76–88. 10.1016/j.neubiorev.2015.03.001.

68. Lo, C.C., and Wang, X.J. (2006). Cortico-basal ganglia circuit mechanism for a decision threshold in reaction time tasks. Nat Neurosci 9, 956–963. 10.1038/nn1722.

69. Fleming, E.A., Field, G.D., Tadross, M.R., and Hull, C. (2024). Local synaptic inhibition mediates cerebellar granule cell pattern separation and enables learned sensorimotor associations. Nat Neurosci 27, 689–701. 10.1038/s41593-023-01565-4.

70. Samsonovich, A., and McNaughton, B.L. (1997). Path integration and cognitive mapping in a continuous attractor neural network model. J Neurosci 17, 5900–5920. 10.1523/JNEUROSCI.17-15-05900.1997.

